# CoCoNuTs: A diverse subclass of Type IV restriction systems predicted to target RNA

**DOI:** 10.1101/2023.07.31.551357

**Authors:** Ryan T. Bell, Harutyun Sahakyan, Kira S. Makarova, Yuri I. Wolf, Eugene V. Koonin

## Abstract

A comprehensive census of McrBC systems, among the most common forms of prokaryotic Type IV restriction systems, followed by phylogenetic analysis, reveals their enormous abundance in diverse prokaryotes and a plethora of genomic associations. We focus on a previously uncharacterized branch, which we denote CoCoNuTs (<u>co</u>iled-<u>co</u>il <u>nu</u>clease tandems) for their salient features: the presence of extensive coiled-coil structures and tandem nucleases. The CoCoNuTs alone show extraordinary variety, with 3 distinct types and multiple subtypes. All CoCoNuTs contain domains predicted to interact with translation system components, such as OB-folds resembling the SmpB protein that binds bacterial transfer-messenger RNA (tmRNA), YTH-like domains that might recognize methylated tmRNA, tRNA, or rRNA, and RNA-binding Hsp70 chaperone homologs, along with RNases, such as HEPN domains, all suggesting that the CoCoNuTs target RNA. Many CoCoNuTs might additionally target DNA, via McrC nuclease homologs. Additional restriction systems, such as Type I RM, BREX, and Druantia Type III, are frequently encoded in the same predicted superoperons. In many of these superoperons, CoCoNuTs are likely regulated by cyclic nucleotides, possibly, RNA fragments with cyclic termini, that bind associated CARF (<u>C</u>RISPR-<u>A</u>ssociated <u>R</u>ossmann <u>F</u>old) domains. We hypothesize that the CoCoNuTs, together with the ancillary restriction factors, employ an echeloned defense strategy analogous to that of Type III CRISPR-Cas systems, in which an immune response eliminating virus DNA and/or RNA is launched first, but then, if it fails, an abortive infection response leading to PCD/dormancy via host RNA cleavage takes over.

## Introduction

All organisms are subject to an incessant barrage of genetic parasites, such as viruses and transposons. Over billions of years, the continuous arms race between hosts and parasites drove the evolution of immense, intricately interconnected networks of diverse defense systems and pathways (Burroughs et al., 2015, Gao et al., 2020, Goldfarb et al., 2015, Bell et al., 2020, Koonin and Aravind, 2002, Swarts et al., 2014). In particular, in the last few years, targeted searches for defense systems in prokaryotes, typically capitalizing on the presence of variable genomic defense islands, have dramatically expanded their known diversity and led to the discovery of a plethora of biological conflict strategies and mechanisms (Gao et al., 2020, Bell et al., 2020, Anantharaman et al., 2012, Kaur et al., 2020).

One of the most ancient and common forms of defense against mobile genetic elements (MGE) is the targeted restriction of nucleic acids. Since the initial discovery of this activity among strains of bacteria resistant to certain viruses, myriad forms of recognition and degradation of nucleic acids have been described, in virtually all life forms. Characterization of the most prominent of these systems, such as restriction-modification (RM), RNA interference (RNAi), and CRISPR-Cas (<u>c</u>lustered <u>r</u>egularly interspaced <u>s</u>hort <u>p</u>alindromic <u>r</u>epeats-<u>C</u>RISPR <u>as</u>sociated genes), has led to the development of a profusion of highly effective experimental and therapeutic techniques, in particular, genome editing and engineering (Loenen et al., 2014, Fire et al., 1998, Agrawal et al., 2003, Makarova et al., 2006, Makarova et al., 2020b, Gasiunas et al., 2012, Jinek et al., 2012).

Historically, the two-component McrBC (<u>m</u>odified <u>c</u>ytosine <u>r</u>estriction) system was the first form of restriction to be described, although the mechanism remained obscure for decades, and for a time, this system was referred to as RglB (<u>r</u>estriction of <u>g</u>lucoseless phages) due to its ability to restrict T-even phage DNA which contained hydroxymethylcytosine, but not glucosylated bases (Raleigh et al., 1989, Luria and Human, 1952, Fleischman et al., 1976, Dila et al., 1990). Today, the prototypical McrBC system, native to *E. coil* K-12, is considered a Type IV (modification-dependent) restriction system that degrades DNA containing methylcytosine (5mC) or hydroxymethylcytosine (5hmC), with a degree of sequence context specificity (Sutherland et al., 1992, Sukackaite et al., 2012).

Type IV restriction enzymes contain at least two components: 1) a dedicated specificity domain that recognizes modified DNA, and 2) an endonuclease domain that cleaves the target (Weigele and Raleigh, 2016, Loenen et al., 2014). In the well-characterized example from *E. coli* K-12, McrB harbors an N-terminal DUF(domain of unknown function)3578 that recognizes methylcytosine. We denote this domain, as its function is not unknown, as ADAM (<u>a d</u>omain with an <u>a</u>ffinity for <u>m</u>ethylcytosine). The ADAM domain is fused to a GTPase domain of the AAA+ ATPase superfamily, the only known GTPase in this clade, which is believed to translocate DNA (Figure 1 A, B, C) (Sutherland et al., 1992, Iyer et al., 2004, Nirwan et al., 2019, Panne et al., 1999). McrC, encoded by a separate gene in the same operon, contains an N-terminal DUF2357 domain, which interacts with the GTPase domain in McrB and stimulates its activity, and is fused to a PD-(D/E)xK superfamily endonuclease (Figure 1 A, B, D) (Niu et al., 2020, Sutherland et al., 1992).

**Figure 1:**
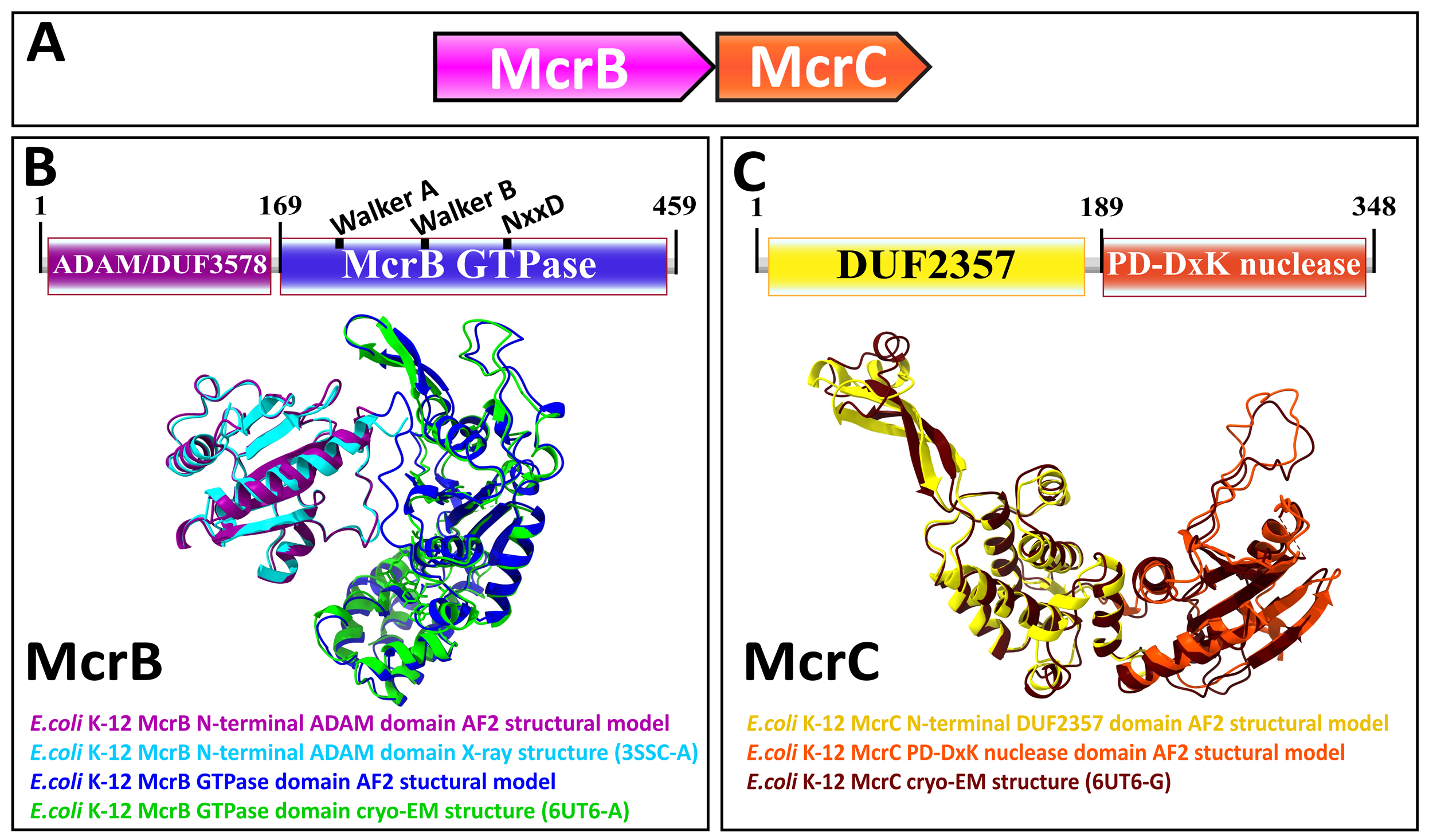
Genetic organization, signature sequence motifs, structural models, and phyletic distribution of McrB GTPases detected in this work. A) McrBC is a two-component restriction system with each component typically (except for extremely rare gene fusions) encoded by a separate gene expressed as a single operon, depicted here and in subsequent figures as arrows pointing in the direction of transcription. In most cases, McrB is the upstream gene in the operon. B) In the prototypical E.coli K-12 McrBC system, McrB contains an N-terminal methylcytosine-binding domain, ADAM/DUF3578, fused to a GTPase of the AAA+ ATPase clade (Sukackaite et al., 2012). This GTPase contains the Walker A and Walker B motifs that are conserved in P-loop NTPases as well as a signature NxxD motif, all of which are required for GTP hydrolysis (Nirwan et al., 2019, Niu et al., 2020, Pieper et al., 1999). An AlphaFold2 structural model of E. coli K-12 ADAM-McrB GTPase fusion protein monomer and separate X-ray diffraction and cryo-EM structures of the ADAM and GTPase domains (Niu et al., 2020, Sukackaite et al., 2012) show a high degree of similarity. C) McrC consists of a PD-DxK nuclease and an N-terminal DUF2357 domain, which comprises a helical bundle with a stalk-like extension that interacts with and activates individual McrB GTPases while they are assembled into hexamers (Niu et al., 2020, Nirwan et al., 2019). An AlphaFold2 structural model and cryo-EM structure of E. coli K-12 McrC monomer with DUF2357-PD-D/ExK architecture (Niu et al., 2020) show a high degree of similarity. The structures were visualized with ChimeraX (Pettersen et al., 2021).

Our recent analysis of the modified base-binding EVE (named for Protein Data Bank (PDB) structural identifier 2eve) domain superfamily demonstrated how the distribution of the EVE-like domains connects the elaborate eukaryotic RNA regulation and RNA interference-related epigenetic silencing pathways to largely uncharacterized prokaryotic antiphage restriction systems (Bell et al., 2020). EVE superfamily domains, which in eukaryotes recognize modified DNA or RNA as part of mRNA maturation or epigenetic silencing functions, are often fused to McrB-like GTPases in prokaryotes, and indeed, these are the most frequently occurring EVE-containing fusion proteins (Bell et al., 2020). These observations motivated us to conduct a comprehensive computational search for McrBC systems, followed by a census of all associated domains, to chart the vast and diverse population of antiviral specificity modules, vital for prokaryotic defense, that also provided important source material during the evolution of central signature features of eukaryotic cells. Here we present the results of this census and describe an extraordinary, not previously appreciated variety of domain architectures of the McrBC family of Type IV restriction systems. In particular, we focus on a major McrBC branch that we denote <u>co</u>iled-<u>co</u>il <u>nu</u>clease <u>t</u>andem (CoCoNuT) systems, which we explore in detail.

## Results and Discussion

### Comprehensive census of McrBC systems

The search for McrBC systems included PSI-BLAST runs against the non-redundant protein sequence database at the NCBI, followed by several filtering strategies (see Methods) to obtain a clean set of nearly 34,000, distributed broadly among prokaryotes. In the subsequent phase of analysis, GTPase domain sequences were extracted from the McrB homolog pool, and DUF2357 and PD-D/ExK nuclease domain sequences were extracted from the McrC homolog pool, leaving as a remainder the fused specificity domains that we intended to classify (although the McrC homologs are not the primary bearers of specificity modules in McrBC systems, they can be fused to various additional domains, including those of the EVE superfamily) (Figure 2, Supplementary Figure S1). Unexpectedly, we found that both the GTPase domains and DUF2357 domains frequently contained insertions into their coding sequences, likely encoding specificity domains and coiled-coils, respectively.

**Figure 2:**
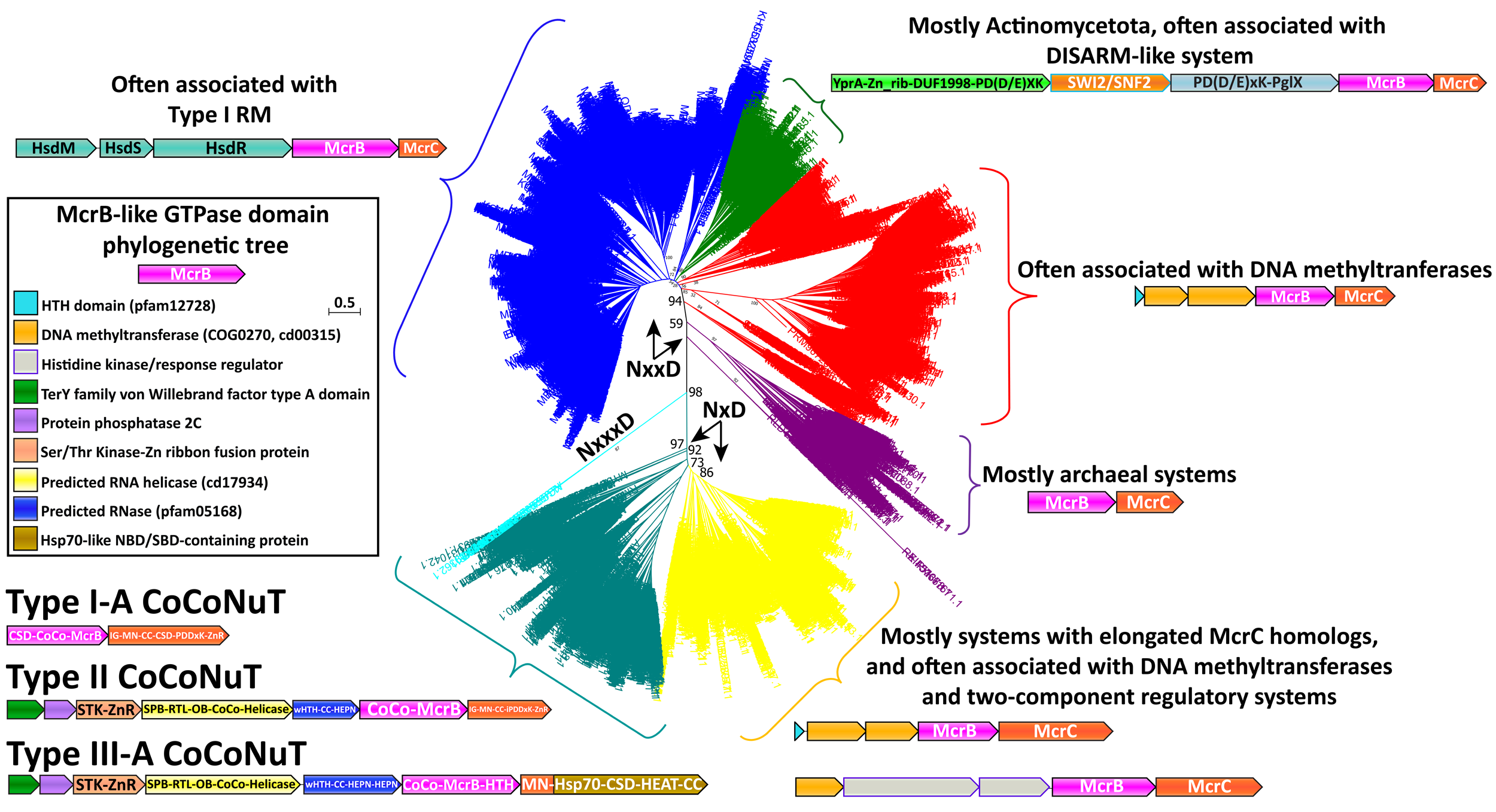
Phylogenetic tree of the McrB-like GTPases. The major clades in the phylogenetic tree of the McrB-like GTPases are distinguished by the distinct versions of the Nx(xx)D signature motif. The teal and yellow groups, with bootstrap support of 97%, have an NxD motif, whereas the blue, green, red, and purple groups, with variable bootstrap support, have an NxxD motif, indicated by the arrows; the sequences in the smaller, cyan clade, with 98% bootstrap support, have an NxxxD motif. Each of the differently colored groups is characterized by distinct conserved genomic associations that are abundant within but not completely confined to the respective groups. This tree was built from the representatives of 90% identity clusters of all validated homologs. Abbreviations of domains: McrB – McrB GTPase domain; CoCo/CC – coiled-coil; MN – McrC N-terminal domain (DUF2357); CSD – cold shock domain; IG – Immunoglobulin (IG)-like beta-sandwich domain; ZnR – zinc ribbon domain; SPB – SmpB-like domain; RTL – RNase toxin-like domain; HEPN – HEPN family nuclease domain; OB – OB-fold domain; iPD-D/ExK – inactivated PD-D/ExK fold; Hsp70-like ATPase – Hsp70-like NBD/SBD; HEAT – HEAT-like helical repeats; YprA – YprA-like helicase domain; DUF1998 - DUF1998 is often found in or associated with helicases and contains four conserved, putatively metal ion-binding cysteine residues; SWI2/SNF2 – SWI2/SNF2-family ATPase; PglX – PglX-like DNA methyltransferase; HsdR/M/S – Type I RM system restriction, methylation, and specificity factors.

Removing variable inserts from the conserved McrBC domains allowed accurate, comprehensive phylogenetic analysis of the McrB GTPase (Figure 2) and McrC DUF2357 (Supplementary Figure S1) families. These two trees were generally topologically concordant and revealed several distinct branches not previously recognized. The branch containing the prototypical McrBC system from *E. coil* K-12 (blue in Figure 2) is characterized by frequent genomic association with Type I RM systems, usually with unidirectional gene orientations and the potential of forming a single operon (Raleigh, 1992). This branch and others, which exhibit two particularly prevalent associations, with a DISARM-like antiphage system (green in Figure 2) (Ofir et al., 2018), and, surprisingly for a Type IV restriction system, with predicted DNA methyltransferases (red in Figure 2), will be the subject of a separate, forthcoming publication.

### A variant of the McrB GTPase signature motif distinguishes a large group of unusual McrBC systems

A major branch (teal and yellow in Figure 2) of the McrBC family is characterized by a conserved deletion within the NxxD GTPase signature motif (where x is any amino acid), found in all McrB-like GTPases, reducing it to NxD (Figure 1 B, Figure 2) (Neuwald et al., 1999, Iyer et al., 2004, Erzberger and Berger, 2006, Niu et al., 2020). The NxD variant of the motif is strictly conserved in these homologs and is usually, but not invariably, followed by a glutamate (E) or a second aspartate (D). No NxD motif McrB GTPase has been characterized, but the extensive study of the NxxD motif offers clues to the potential impact of the motif shortening. The asparagine (N) residue is strictly required for GTP binding and hydrolysis (Pieper et al., 1999). As this residue is analogous to sensor-1 in ATP-hydrolyzing members of the AAA+ family, it can be predicted to position a catalytic water molecule for nucleophilic attack on the γ-phosphate of an NTP (Erzberger and Berger, 2006, Colicelli, 2004, Bourne et al., 1991, Niu et al., 2020, Pieper et al., 1999). A recent structural analysis has shown that the aspartate interacts with a conserved arginine/lysine residue in McrC which, via a hydrogen-bonding network, resituates the NxxD motif in relation to its interface with the Walker B motif such that, together, they optimally position a catalytic water to stimulate hydrolysis (Niu et al., 2020). Accordingly, the truncation of this motif might be expected to modulate the rate of hydrolysis, and potentially compel functional association with only a subset of McrC homologs containing compensatory mutations. The NxD branch is characterized by many McrC homologs with unusual features, such as predicted RNA-binding domains, that might not be compatible with conventional McrB GTPases from the NxxD clade, which often occur in the same genomes (see below).

Many of these NxD GTPases are contextually associated with DNA methyltransferases, like the NxxD GTPases, but are distinguished by additional complexity in the domains fused to the McrC homologs and the frequent presence of two-component regulatory system genes in the same operon (yellow in Figure 2). We also detected a small number of GTPases with an insertion in the signature motif (cyan in Figure 2), expanding it to NxxxD, which are methyltransferase-associated as well. This association is likely to be ancestral, as it is found in all 3 branches of the McrB GTPase tree with different signature motif variations.

The most notable feature of the NxD branch of GTPases is a large clade characterized by fusion to long coiled-coil domains (teal in Figure 2, Supplementary Table S1), a derived and distinctly different architecture from the small, modular DNA-binding specificity domains typically found in canonical McrB homologs. The McrC homologs associated with these coiled-coil McrB GTPase fusions also often lack the PD-D/ExK endonuclease entirely or contain a region AlphaFold2 (AF2) predicted to adopt the PD-(D/E)xK endonuclease fold, but with inactivating replacements of catalytic residues (Supplementary Table S1). However, in the cases where the McrC nuclease is missing or likely inactive, these systems are always encoded in close association, often with overlapping reading frames, with HEPN (<u>h</u>igher <u>e</u>ukaryotes and <u>p</u>rokaryotes <u>n</u>ucleotide-binding) ribonuclease domains, or other predicted nucleases (Figure 3, Supplementary Figure S2) (Pillon et al., 2021). Based on these features of domain architecture and genomic context, we denote these uncharacterized McrBC-containing operons <u>co</u>iled-<u>co</u>il <u>nu</u>clease tandem (CoCoNuT) systems.

**Figure 3:**
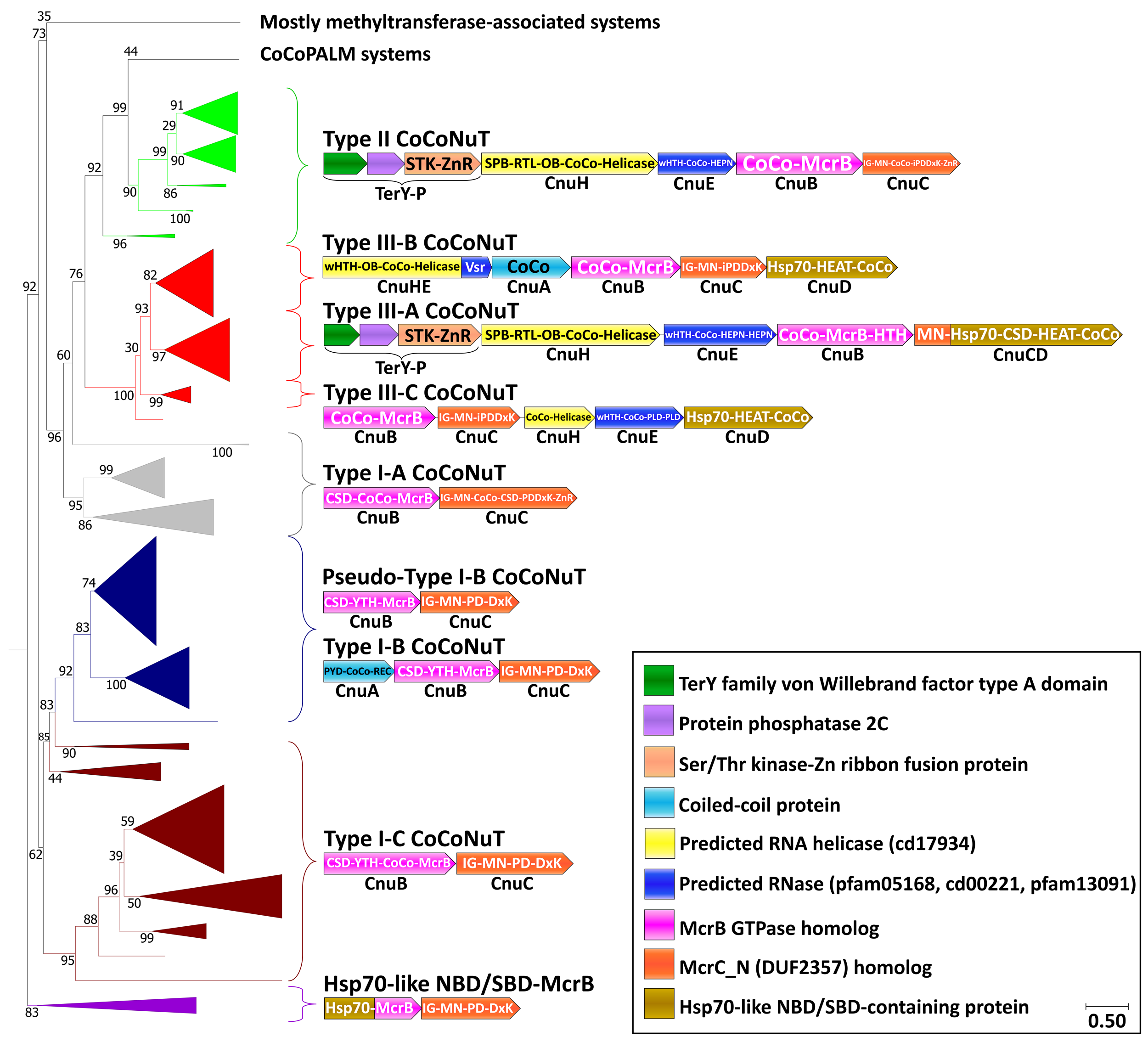
CoCoNuT system phylogeny and classification. The figure shows the detailed phylogeny of McrB GTPases from CoCoNuT systems and their close relatives. All these GTPases possess an NxD GTPase motif rather than NxxD. CoCoNuT systems are abundant in Pseudomonadota and Terrabacteria, with the various subtypes generally concentrated in a more restricted range of species, as indicated. This tree was built from the representatives of 90% clustering of all validated homologs. Abbreviations of domains: McrB – McrB GTPase domain; CoCo – coiled-coil; MN – McrC N-terminal domain (DUF2357); CSD – cold shock domain; YTH – YTH-like domain; IG – Immunoglobulin (IG)-like beta-sandwich domain; Hsp70 – Hsp70-like NBD/SBD; HEAT – HEAT-like helical repeats; ZnR – zinc ribbon domain; PYD – pyrin/CARD-like domain; SPB – SmpB-like domain; RTL – RNase toxin-like domain; HEPN – HEPN family nuclease domain; OB – OB-fold domain; iPD-D/ExK – inactivated PD-D/ExK fold; REC – Phosphoacceptor receiver-like domain; PLD – Phospholipase D-like nuclease domain; Vsr – very-short-patch-repair PD-D/ExK nuclease-like domain. Underneath each gene is a proposed protein name, with Cnu as an abbreviation for <u>C</u>oCo<u>Nu</u>T.

The deepest branching group of the NxD GTPase systems is typified by a fusion of an Hsp70-like ATPase nucleotide-binding domain and substrate-binding domain (NBD/SBD) to the McrB GTPase domain (purple in Figure 3, Supplementary Figure S2, Supplementary Table S1). Hsp70 is an ATP-dependent protein chaperone that binds exposed hydrophobic peptides and facilitates protein folding (Mayer, 2021). It also associates with AU-rich mRNA, and in some cases, such as the bacterial homolog DnaK, C-rich RNA, an interaction involving both the NBD and SBD (Kishor et al., 2017, Zimmer et al., 2001, Kishor et al., 2013). These systems might be functionally related to the CoCoNuTs, given that an analogous unit is encoded by a type of CoCoNuT system where similar domains are fused to the McrC homolog rather than to the McrB GTPase homolog (see below). In another large clade of CoCoNuT-like systems, the GTPase is fused to a domain homologous to FtsB, an essential bacterial cell division protein containing transmembrane and coiled-coil helices (Supplementary Figure S2, Supplementary Table S1) (Khadria and Senes, 2013). They are associated with signal peptidase family proteins likely to function as pilus assembly factors (Supplementary Table S1) (Colicelli, 2004) and usually contain coiled-coil domains fused to both the McrB and McrC homologs. Therefore, we denote them <u>co</u>iled-<u>co</u>il and <u>p</u>ilus <u>a</u>ssembly linked to <u>M</u>crBC (CoCoPALM) systems (Supplementary Figure S2).

Here, we focus on the CoCoNuTs, whereas the CoCoPALMs and the rest of the NxD GTPase methyltransferase-associated homologs will be explored in a separate, forthcoming publication. CoCoNuT systems are extremely diverse and represented in a wide variety of bacteria, particularly Pseudomonadota and Bacillota, but are nearly absent in archaea (Figure 4).

**Figure 4:**
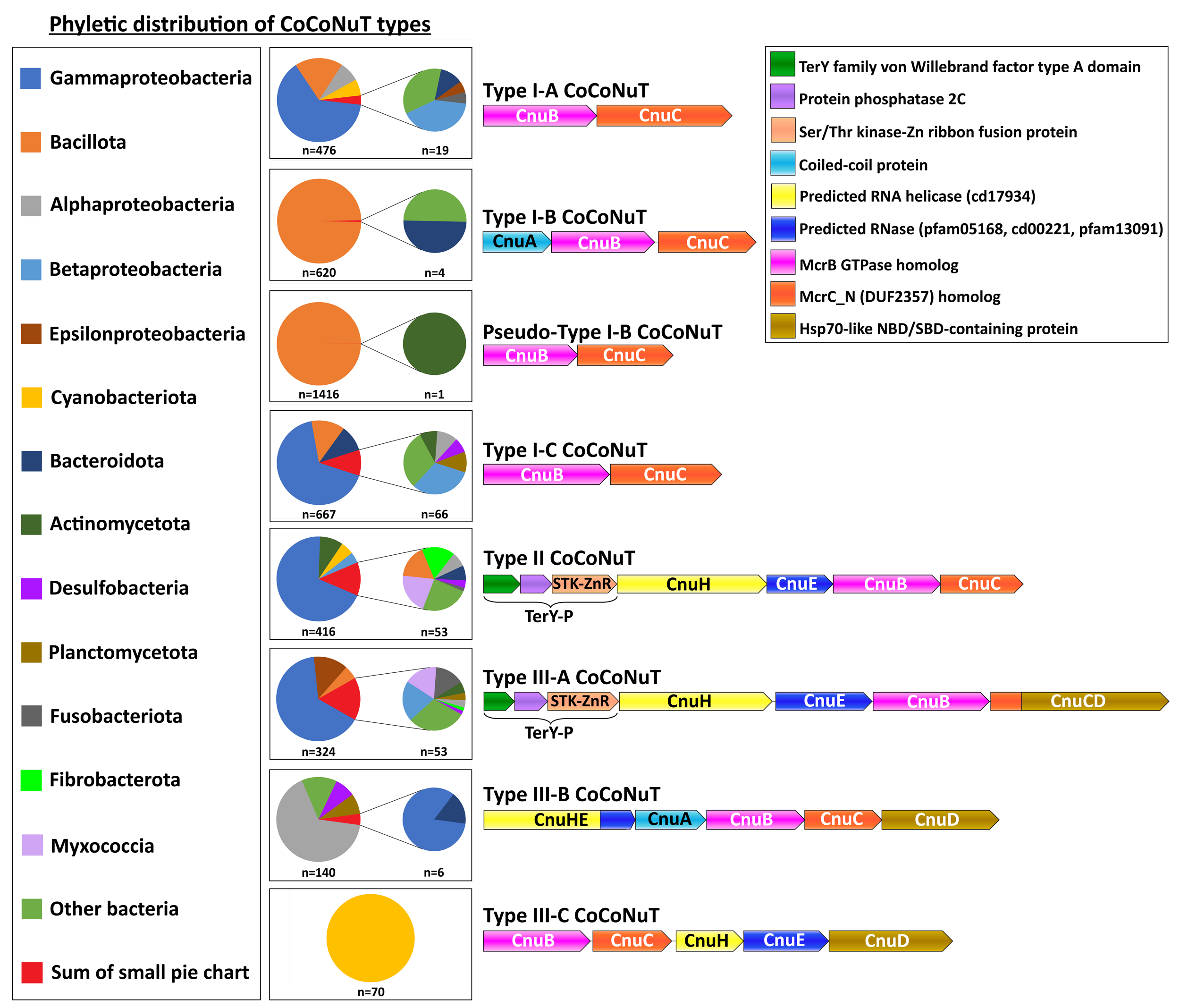
Phyletic distribution of CoCoNuTs. The phyletic distribution of CnuB/McrB GTPases in CoCoNuT systems found in genomic islands with distinct domain compositions. Most CoCoNuTs are found in either Bacillota or Pseudomonadota, with particular abundance in Gammaproteobacteria. Type I-B and the related Pseudo-Type I-B CoCoNuTs are restricted mainly to Bacillota. In contrast, the other types are more common in Pseudomonadota, but can be found in a wide variety of bacteria. Type III-B and III-C are primarily found in Alphaproteobacteria and Cyanobacteriota, respectively.

### Type I CoCoNuT systems

We classified the CoCoNuTs into 3 types and 7 subtypes based on the GTPase domain phylogeny and conserved genomic context (Figure 3). Type I-A and Type I-C systems consist of McrB and McrC homologs only, which we denote CnuB and CnuC (Figure 3, Figure 5, Supplementary Figure S7, Supplementary Figure S9). Type I-A is distinguished from all other CoCoNuT types by a helical insert into the CnuB GTPase domain, between the Walker B and NxD motifs (Figure 5, Supplementary Figure S7).

**Figure 5:**
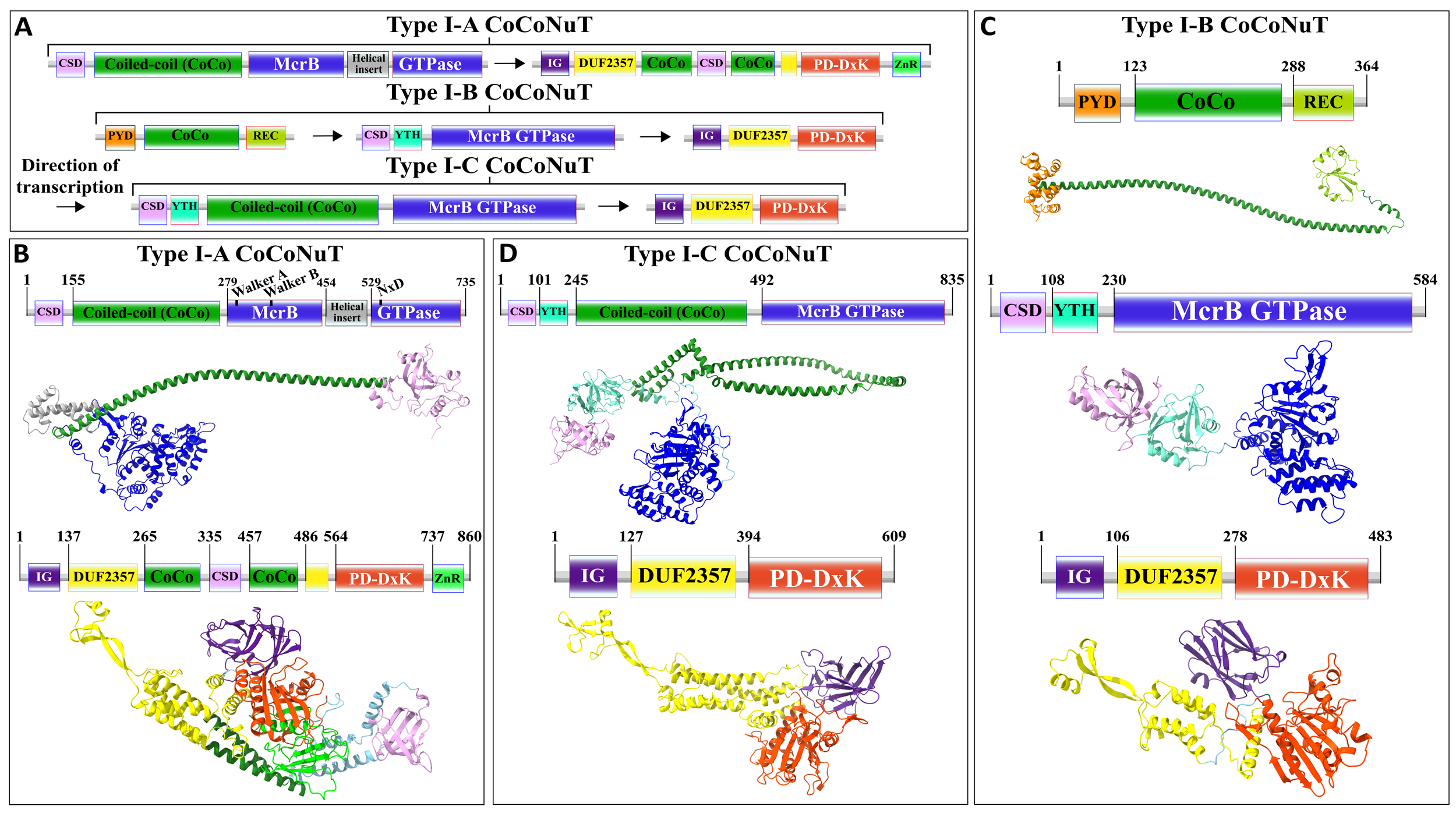
Domain composition, operon organization, and AlphaFold2 structural predictions of components of the Type I CoCoNuT systems. A) Type I CoCoNuT domain composition and operon organization. The arrows indicate the direction of transcription. B-D) High quality (Average pLDDT > 80), representative AlphaFold2 structural predictions for protein monomers in B) Type I-A CoCoNuT systems (CnuB and CnuC, from top to bottom), C) Type I-B (CnuA, CnuB, and CnuC, from top to bottom), and D) Type I-C CoCoNuT systems (CnuB and CnuC, from top to bottom). Models were generated from representative sequences with the following GenBank accessions (see Supplementary Data for sequences and locus tags): ROR86958.1 (Type I-A CnuB), APL73566.1 (Type I-A CnuC), TKH01449.1 (Type I-B CnuA), GED20858.1 (Type I-B CnuB), GED20857.1 (Type I-B CnuC), GFD85286.1 (Type I-C CnuB), MBV0932851.1 (Type I-C CnuC). Abbreviations of domains: CSD – cold shock domain; YTH – YTH-like domain; CoCo – coiled-coil; IG – Immunoglobulin (IG)-like beta-sandwich domain; ZnR – zinc ribbon domain; PYD – pyrin/CARD-like domain; REC – Phosphoacceptor receiver-like domain. These structures were visualized with ChimeraX (Pettersen et al., 2021).

Type I-B systems usually encode a separate coiled-coil protein, which we denote CnuA, in addition to CnuB/McrB and CnuC/McrC, with no coiled-coil fused to the McrB-like GTPase domain in CnuB (Figure 3, Figure 5, Supplementary Figure S8). CnuA is fused at the N-terminus to a pyrin (PYD)/CARD (<u>c</u>aspase <u>a</u>ctivation and <u>r</u>ecruitment <u>d</u>omain)-like helical domain and at the C-terminus to a phosphoacceptor receiver (REC) domain (Figure 5, Supplementary Table S1). The association of the PYD/CARD-like domains with CoCoNuTs suggests involvement in a programmed cell death (PCD)/abortive infection-type response, as they belong to the DEATH domain superfamily and are best characterized in the context of innate immunity, inflammasome formation, and PCD (Park et al., 2007). The REC domains constitute one of the components of two-component regulatory systems. They are targeted for phosphorylation by histidine kinases (Stock et al., 2000), which could be a mechanism of Type I-B CoCoNuT regulation.

In contrast to Type II and Type III CnuB homologs, which contain only coiled-coils fused at their N-termini (Figure 6), cold-shock domain (CSD)-like OB-folds are usually fused at the N-termini of Type I CoCoNuT CnuB/McrB proteins, in addition to the coiled-coils (except for the majority of Type I-B, where the coiled-coils are encoded separately) (Figure 5, Supplementary Table S1) (Amir et al., 2018). Often, in Type I-B and I-C, but not Type I-A, a YTH-like domain, a member of the modified base-binding EVE superfamily, is present in CnuB as well, between the CSD and coiled-coil, or in Type I-B, between the CSD and the GTPase domain (Figure 5, Supplementary Table S1) (Bell et al., 2020, Hazra et al., 2019, Liao et al., 2018).

**Figure 6:**
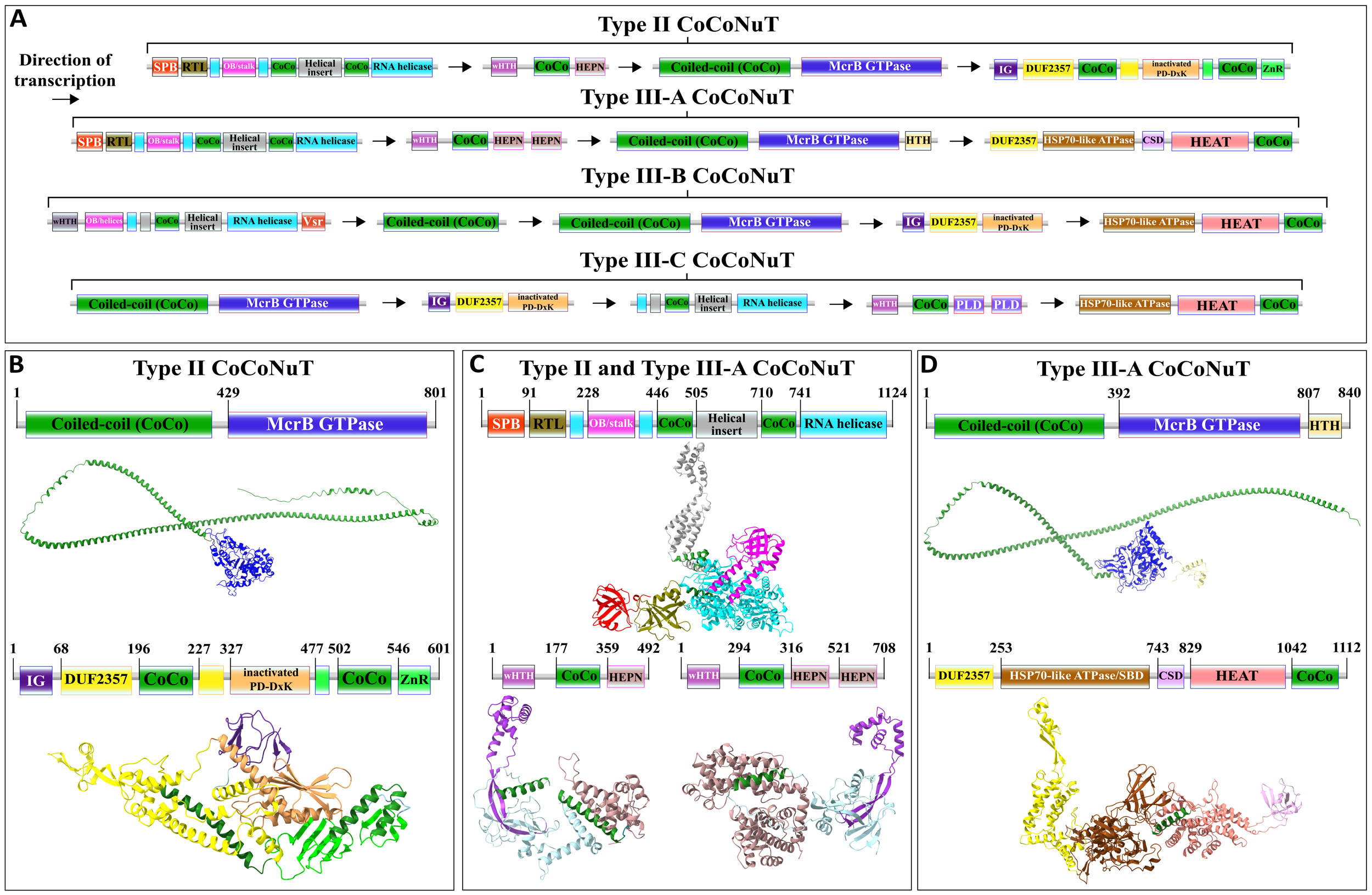
Domain composition, operon organization, and AlphaFold2 structural predictions for core protein components of Type II and III CoCoNuT systems. A) Type II and III CoCoNuT domain composition and operon organization. The arrows indicate the direction of transcription. Type II and III-A systems very frequently contain TerY-P systems as well, but not invariably, and these are never found in Type III-B or III-C, thus, we do not consider them core components. B-D) High quality (Average pLDDT > 80), representative AlphaFold2 structural predictions for protein monomers in B) Type II CoCoNuT systems (CnuB and CnuC, from top to bottom), C) Type II and III-A CoCoNuT systems (CnuH at the top, Type II CnuE on the bottom left, Type III-A CnuE on the bottom right), and D) Type III-A CoCoNuT systems (CnuB and CnuC, from top to bottom). Models were generated from representative sequences with the following GenBank accessions (see Supplementary Data for sequences and locus tags): AMO81401.1 (Type II CnuB), AVE71177.1 (Type II CnuC), AMO81399.1 (Type II and III-A CoCoNuT CnuH), AVE71179.1 (Type II CnuE), ATV59464.1 (Type III-A CnuE), PNG83940.1 (Type III-A CnuB), NMY00740.1 (Type III-A CnuC). Abbreviations of domains: CSD – cold shock domain; CoCo – coiled-coil; IG – Immunoglobulin (IG)-like beta-sandwich domain; ZnR – zinc ribbon domain; SPB – SmpB-like domain; RTL – RNase toxin-like domain; HEPN – HEPN family nuclease domain; OB/stalk – OB-fold domain attached to a helical stalk-like extension of ATPase; Vsr – very-short-patch-repair PD-D/ExK nuclease-like domain; PLD – Phospholipase D family nuclease domain; HEAT – HEAT-like helical repeats. These structures were visualized with ChimeraX (Pettersen et al., 2021).

The CnuC/McrC proteins in Type I CoCoNuTs, as well as those in Type II and Type III-B and III-C, all contain an immunoglobulin-like N-terminal beta-sandwich domain of unknown function, not present in the *E. coli* K-12 McrC homolog, similar to a wide range of folds from this superfamily with diverse roles (Figure 5, Supplementary Table S1) (Natarajan et al., 2015, Halaby et al., 1999). It is also present in non-CoCoNuT McrC homologs associated with McrB GTPase homologs in the NxD clade, implying its function is not specific to the CoCoNuTs. In the Type I-B and I-C CoCoNuT CnuC homologs, these domains most closely resemble Rho GDP-dissociation inhibitor 1, suggesting that they may be involved in the regulation of CnuB/McrB GTPase activity (Dovas and Couchman, 2005).

We also detected a close relative of Type I-B CoCoNuT in many *Bacillus* species, which we denoted Pseudo-Type I-B CoCoNuT because it lost the separate coiled-coil protein CnuA. Pseudo-Type I-B CoCoNuT occurs in genomic contexts suggesting a role in overcrowding-induced stress responses (Figure 3, Supplementary Figure S4, Supplementary Note S1).

### Type II and III CoCoNuT systems

Type II and III CoCoNuT CnuB/McrB GTPase domains branch from within Type I-A and are encoded in a nearly completely conserved genomic association with a Superfamily 1 (SF1) helicase of the UPF1-like clade, which we denote CnuH (Figure 3, Supplementary Table S1) (Gorbalenya and Koonin, 1993, Fairman-Williams et al., 2010). The UPF1-like family encompasses helicases with diverse functions acting on RNA and single-stranded DNA (ssDNA) substrates, and notably, the prototypical UPF1 RNA helicase and its closest relatives are highly conserved in eukaryotes, where they play a critical role in the nonsense-mediated decay (NMD) RNA surveillance pathway (Cheng et al., 2007, Chakrabarti et al., 2011).

SF1 helicases are composed of two RecA-like domains which together harbor a series of signature motifs required for the ATPase and helicase activities, including the Walker A and Walker B motifs conserved in P-loop NTPases, which are located in the N-terminal RecA-like domain (Fairman-Williams et al., 2010). In all 4 Type II and Type III CoCoNuT subtypes, following the Walker A motif, the CnuH helicases contain a large helical insertion, with some of the helices predicted to form coiled-coils (Figure 6). The Type II, Type III-A, and Type III-B CoCoNuT CnuH helicases contain an OB-fold domain that, in Type II and Type III-A, is flanked by helices predicted to form a stalk-like helical extension of the N-terminal RecA-like domain, a structural feature characteristic of the entire UPF1/DNA2-like helicase family within SF1 (Chakrabarti et al., 2011, Zhou et al., 2015, Kalathiya et al., 2019) (Figure 6). DALI comparisons show that the CnuH predicted OB-fold domain in Type II CoCoNuTs is similar to the OB-fold domain in UPF1 and related RNA helicases SMUBP-2 and SEN1, and this holds for Type III-A as well, although, in these systems, the best DALI hits are to translation factor components such as EF-Tu domain II (Supplementary Table S1) (Chakrabarti et al., 2011, Morse et al., 2020). The Type III-B CnuH predicted OB-folds also match that of UPF1, albeit with lower statistical support (Supplementary Table S1).

In Type II and Type III-A CoCoNuTs, CnuH is fused at the N-terminus to a second OB-fold domain similar to that of SmpB (<u>sm</u>all <u>p</u>rotein B) (Figure 6, Supplementary Figure S5, Supplementary Table S1), which we denote SPB (<u>S</u>m<u>pB</u>-like). SmpB binds to SsrA RNA, also known as transfer-messenger RNA (tmRNA), and is required for tmRNA to rescue stalled ribosomes, via entry into their A-sites with its alanine-charged tRNA-like domain (Barends et al., 2001, Himeno et al., 2014, Guyomar et al., 2021). The SPB domain is around 20 amino acids shorter on average than SmpB itself, and a helix that is conserved in SmpB orthologs and interacts with the tmRNA is absent in the CoCoNuT OB folds (Supplementary Figure S5) (Bessho et al., 2007). However, two other structural elements of SmpB involved in binding tmRNA are present (Supplementary Figure S5) (Gutmann et al., 2003). The N-terminal OB-folds in Type II and III-A CnuH homologs also resemble prokaryotic HIRAN domains, which are uncharacterized, but have eukaryotic homologs fused to helicases that bind ssDNA (Supplementary Figure S5) (Chavez et al., 2018). These HIRAN domains, however, do not overlay with the CoCoNuT domains any better than SmpB (Supplementary Figure S5), lack a small helix that SmpB and the CoCoNuT domains share, and likely being DNA-binding, do not fit the other pieces of evidence we gathered that all suggest an RNA-binding role for this domain in the CoCoNuTs (see below).

The Type II and Type III-A CoCoNuT CnuH helicases contain an additional domain, structurally similar to the RelE/Colicin D RNase fold (Gucinski et al., 2019), which we denote <u>R</u>Nase toxin-like (RTL), located between the SPB OB-fold and the helicase (Figure 6, Supplementary Table S1). The proteins of this family are ribosome-dependent toxins that cleave either mRNA or tRNA in the ribosomal A-site (Pedersen et al., 2003). This domain was identified by structural similarity search with the AF2 models, but no sequence conservation with characterized members of this family was detected, leaving it uncertain whether the RTL domain is an active nuclease. However, considerable divergence in sequence is not unusual in this toxin family (Guglielmini and Van Melderen, 2011, Goeders et al., 2013). In Type III-B CoCoNuTs, a wHTH domain resembling the archaeal ssDNA-binding protein Sul7s is fused to the CnuH N-termini (Figure 6, Supplementary Table S1).

We searched for additional homologs of CnuH (see Methods). Our observations of the contextual associations, both of the CoCoNuTs and their relatives, showed that helicases of this large family are typically encoded in operons with downstream genes coding for an elongated wHTH domain fused at its C-terminus to a variety of effectors, generally, nucleases, which we denote CnuE (Figure 3, Figure 6). In the CoCoNuTs, except for Types III-B and III-C, these CnuE effectors are HEPN ribonucleases, with one HEPN domain in Type II and two in Type III-A (Figure 6, Supplementary Table S1). In Type III-B, the effector is a Vsr (<u>v</u>ery-<u>s</u>hort-patch <u>r</u>epair)-like PD-(D/E)xK family endonuclease (Tsutakawa et al., 1999) fused directly to the helicase, with no wHTH domain present, whereas in Type III-C, a distorted version of the elongated wHTH domain is fused to two Phospholipase D (PLD) family endonuclease domains (Figure 6, Supplementary Table S1). All these nucleases can degrade RNA, and some, such as HEPN, have been found to cleave RNA exclusively (Pillon et al., 2021, Ipsaro et al., 2012, Mendez et al., 2018, Laganeckas et al., 2011, Songailiene et al., 2020). We also detected coiled-coils in the region between the wHTH and effector domains in Type II, Type III-A, and Type III-C CoCoNuT CnuE homologs. Multimer structural modeling with AlphaFold2 suggests that the wHTH domain in CnuE might interact with the coiled-coil-containing helical insertion of CnuH, perhaps mediated by the coiled-coils in each protein, to couple the ATP-driven helicase activity to the various nuclease effectors (Supplementary Figure S11). The accuracy of this model notwithstanding, the fusion of the Vsr-like effector to CnuH in Type III-B CoCoNuTs and the similarity of this system to the other CoCoNuT types strongly suggests that the CnuE effector proteins in these systems form complexes with their respective CnuH helicases. An additional factor in potential complexing by these proteins is the presence of coiled-coils in the associated CnuB/McrB and CnuC/McrC homologs, which may interact with the coiled-coils in CnuH and CnuE. Type III-B CoCoNuTs also code for a separate coiled-coil protein, which we denote CnuA, as it resembles the CnuA protein encoded in Type I-B CoCoNuTs (Figure 3, Figure 5, Figure 6). However, it is distinguished from Type I-B CnuA in containing no recognizable domains other than the coiled-coil (Figure 3, Figure 5, Figure 6). This also could potentially interact with other coiled-coil proteins in the system.

Type II and Type III-A CoCoNuTs, the most widespread varieties apart from Type I, also include conserved genes coding for a “TerY-P” triad. TerY-P consists of a TerY-like von Willebrand factor type A (VWA) domain, a protein phosphatase 2C-like enzyme, and a serine/threonine kinase (STK) fused at the C-terminus to a zinc ribbon (Figure 3, Figure 6, Supplementary Table S1). TerY-P triads are involved in tellurite resistance, associated with various predicted DNA restriction and processing systems, and are predicted to function as a metal-sensing phosphorylation-dependent signaling switch (Anantharaman et al., 2012). In addition, TerY-P-like modules, in which the kinase is fused at the C-terminus to an OB-fold rather than a zinc ribbon, have been recently shown to function as stand-alone antiphage defense systems (Gao et al., 2020). The OB-fold fusion suggests that this kinase interacts with an oligonucleotide and raises the possibility that the zinc ribbon, which occupies the same position in the CoCoNuTs, is also nucleic acid-binding. Almost all CoCoNuT systems containing *cnuHE* operons also encompass TerY-P, with a few exceptions among Terrabacterial Type II systems and Myxococcal Type III-A systems, implying an important contribution to their function (Supplementary Table S3). However, the complete absence of the TerY-P module in Type III-B and Type III-C systems implies that when different nucleases and other putative effectors fused to the helicase are present, TerY-P is dispensable for the CoCoNuT activity. Therefore, we do not consider them to be core components of these systems.

Type II and Type III-A CoCoNuTs have similar domain compositions, but a more detailed comparison reveals substantial differences. The CnuE proteins in Type II contain one HEPN domain with the typical RxxxxH RNase motif conserved in most cases, whereas those in Type III-A contain two HEPN domains, one with the RxxxxH motif, and the other, closest to the C-terminus, with a shortened RxH motif. Furthermore, Type III-A CnuB/McrB GTPase homologs often contain C-terminal HTH-domain fusions absent in Type II (Figure 6). Finally, striking divergence has occurred between the Type II and III-A CnuC/McrC homologs. Type II CnuCs resemble Type I-A CnuCs, which contain N-terminal immunoglobulin-like beta-sandwich domains, PD-D/ExK nucleases, zinc ribbon domains, and insertions into DUF2357 containing coiled-coils and a CSD-like OBD. However, in Type II CnuCs, this domain architecture underwent reductive evolution (Figure 5, Figure 6, Supplementary Table S1). In particular, the beta-sandwich domain and coiled-coils are shorter, the CSD was lost, the nuclease domain was inactivated, and in many cases, the number of Zn-binding CPxC motifs was reduced from three to two (Figure 5, Figure 6, Supplementary Table S1). This degeneration pattern could indicate functional replacement by the associated CnuH and CnuE proteins, often encoded in reading frames overlapping with the start of the *cnuBC* operon.

By contrast, Type III-A CnuC/McrC homologs entirely lost the beta-sandwich domain, PD-D/ExK nuclease, and zinc ribbon found in Type I-A and Type II, but gained an Hsp70-like NBD/SBD unit similar to those that are fused to the McrB GTPase in early branching members of the NxD clade. They have also acquired a helical domain similar to the HEAT repeat family, and in some cases, a second CSD (Figure 3, Figure 5, Figure 6, Supplementary Table S1). Most Type III-A CnuCs contain a CSD and coiled-coils, and thus, resemble Type I-A, but the positioning of these domains, which in Type I-A are inserted into the DUF2357 helix bundle, is not conserved in Type III-A, where these domains are located outside DUF2357 (Figure 5, Figure 6).

Hsp70-like ATPase NBD/SBD domains and HEAT-like repeats are fused to the CnuC/McrC N-terminal DUF2357 domain in Type III-A CoCoNuT, but their homologs in Type III-B and III-C are encoded by a separate gene. We denote these proteins CnuD and CnuCD, the latter for the CnuC-CnuD fusions in Type III-A. The separation of these domains in Type III-B and III-C implies that fusion is not required for their functional interaction with the CnuBC/McrBC systems (Figure 6 A, Supplementary Table S1). CnuD proteins associated with both Type III-B and III-C contain predicted coiled-coils, suggesting that they might interact with the large coiled-coil in the CnuB homologs (Figure 6 A).

In the CnuD homologs found in the CoCoNuTs and fused to NxD McrB GTPases, Walker B-like motifs (Yamamoto et al., 2014) are usually, but not invariably, conserved, whereas the sequences of the helical domains adjacent to the motifs are more strongly constrained. Walker A-like motifs (Chang et al., 2008) are present but degenerate (Supplementary Figure S13). Therefore, it appears likely that the CoCoNuT CnuD homologs bind ATP/ADP but hydrolyze ATP with extremely low efficiency, at best. Such properties in an Hsp70-like domain are better compatible with RNA binding than unfolded protein binding or remodeling, suggesting that these CnuD homologs may target the respective systems to aberrant RNA. Many Hsp70 homologs have been reported to associate with ribosomes (Mayer, 2021, Willmund et al., 2013), which could also be true for the CoCoNuT Cnu(C)Ds.

Consistent with this prediction, Type II and III-A CoCoNuTs likely target RNA rather than DNA, given that the respective operons encode HEPN RNases, typically the only recognizable nuclease in these systems. All CnuC/McrC homologs in Type II or III CoCoNuT systems lack the PD-D/ExK catalytic motif that is required for nuclease activity, although for Type II and Type III-B and III-C, but not Type III-A, structural modeling indicates that the inactivated nuclease domain was retained, likely for a nucleic acid-binding role (Figure 6, Supplementary Table S1). In Type

III-A CoCoNuT, the nuclease domain was lost entirely and replaced by the Hsp70-like NBD/SBD domain with RNA-binding potential described above (Figure 6, Supplementary Table S1). Often, one or two CSDs, generally RNA-binding domains, although capable of binding ssDNA, are fused to Type III-A CnuC homologs as well (Figure 6, Supplementary Table S1) (Heinemann and Roske, 2021). RNA targeting capability of Type III-B and III-C can perhaps be inferred from their similarity to Type III-A in encoding CnuD Hsp70-like proteins. Moreover, higher-order associations of Type II and Type III-A CoCoNuT systems with various DNA restriction systems suggest a two-pronged DNA and RNA restriction strategy reminiscent of Type III CRISPR-Cas (see below).

We suspect that RNA targeting is an ancestral feature of the CoCoNuT systems. Several observations are compatible with this hypothesis:

1. Most of the CoCoNuts encompass HEPN nucleases that appear to possess exclusive specificity for RNA.
2. CSD-like OB-folds are pervasive in these systems, being present in the CnuB/McrB homologs of all Type I subtypes and in Type I-A and Type III-A CnuC/McrC homologs. As previously noted, these domains typically bind RNA, although they could bind ssDNA as well.
3. YTH-like domains are present in most Type I CoCoNuts, particularly, in almost all early branching Type I-B and I-C systems, suggesting that the common ancestor of the CoCoNuTs contained such a domain. YTH domains in eukaryotes sense internal N6-methyladenosine (m6A) in mRNA (Hazra et al., 2019, Liao et al., 2018, Patil et al., 2018).
4. Type II and Type III-B/III-C CoCoNuTs, which likely target RNA, given the presence of HEPN domains and Hsp70 NBD/SBD homologs, retain inactivated PD-DxK nuclease domains, suggesting that these domains contribute an affinity for RNA inherited from Type I-A CoCoNuTs. PD-(D/E)xK nucleases are generally DNA-specific, however, some examples of RNase activity have been reported (Mendez et al., 2018, Laganeckas et al., 2011). The inactivated PD-DxK domains might also bind DNA from which the target RNA is transcribed.

### Complex higher-order associations between CoCoNuTs, CARF domains, and other defense systems

Genomic neighborhoods of many Type II and Type III-A CoCoNuTs encompass complex operonic associations with genes encoding several types of CARF (<u>C</u>RISPR-<u>A</u>ssociated <u>R</u>ossmann <u>F</u>old) domain-containing proteins. This connection suggests multifarious regulation by cyclic (oligo)nucleotide second messengers synthesized in response to viral infection and bound by CARF domains (Makarova et al., 2020a, McMahon et al., 2020, Zhu et al., 2021). Activation of an effector, most often a nuclease, such as HEPN or PD-(D/E)xK, by a CARF bound to a cyclic (oligo)nucleotide is a crucial mechanism of CBASS (<u>c</u>yclic oligonucleotide-<u>b</u>ased <u>a</u>ntiphage <u>s</u>ignaling <u>s</u>ystem) as well as Type III CRISPR-Cas systems (McMahon et al., 2020, Makarova et al., 2020a, Zhu et al., 2021). These CARF-regulated enzymes generally function as a fail-safe that eventually induces PCD/dormancy when other antiphage defenses fail to bring the infection under control and are deactivated, typically through cleavage of the second messenger by a RING nuclease, if other mechanisms succeed (Makarova et al., 2012, Koonin and Zhang, 2017, Makarova et al., 2020a, Koonin and Krupovic, 2019).

The presence of CARFs could implicate the HEPN domains of these systems as PCD effectors that would carry out non-specific RNA degradation in response to infection. Surprisingly, however, most of these CARF domain-containing proteins showed the highest similarity to RtcR, a sigma54 transcriptional coactivator of the RNA repair system RtcAB with a CARF-ATPase-HTH domain architecture, suggesting an alternative functional prediction (Supplementary Figure S14, Supplementary Table S1). Specifically, by analogy with RtcR, CoCoNuT-associated CARF domain-containing proteins might bind (t)RNA fragments with 2’,3’ cyclic phosphate ends (Kotta-Loizou et al., 2022, Hughes et al., 2020). This interaction could promote transcription of downstream genes, in this case, genes encoding CoCoNuT components, through binding an upstream activating sequence by the HTH domain fused to the CARF-ATPase C-terminus (Hughes et al., 2020, Kotta-Loizou et al., 2022).

The manifold biological effects of tRNA-like fragments are only beginning to be appreciated. Lately, it has been shown that bacterial anticodon nucleases, in response to infection and DNA degradation by phages, generate tRNA fragments, likely a signal of infection and a defensive strategy to slow down the translation of viral mRNA, and that phages can deploy tRNA repair enzymes and other strategies to counteract this defense mechanism (Bitton et al., 2015, van den Berg et al., 2023, Kaufmann, 2000, Ishita et al., 2021). Moreover, the activity of the HEPN ribonucleases in the CoCoNuTs themselves would produce RNA cleavage products with cyclic phosphate ends that might be bound by the associated RtcR-like CARFs (Shigematsu et al., 2018, Pillon et al., 2021), in a potential feedback loop.

Such a CoCoNuT mechanism could complement the function of RtcAB, as RtcA is an RNA cyclase that converts 3L-phosphate RNA termini to 2L,3L-cyclic phosphate and thus, in a feedback loop, generates 2’,3’-cyclic phosphate RNA fragments that induce expression of the operon (Genschik et al., 1998, Das and Shuman, 2013, Hughes et al., 2020). Acting downstream of RtcA, RtcB is an RNA ligase that joins 2’,3’-cyclic phosphate RNA termini to 5’-OH RNA fragments, generating a 5’-3’ bond and, in many cases, reconstituting a functional tRNA (Tanaka and Shuman, 2011). Recent work has shown that, in bacteria, the most frequent target of RtcB is SsrA, the tmRNA (Kotta-Loizou et al., 2022). Intriguingly, as described above, in order to rescue stalled ribosomes, tmRNA must bind to SmpB, an OB-fold protein highly similar to the predicted structures of the SPB domains fused at the N-termini of the CnuH helicases in Type II and III-A CoCoNuTs, which are the only types that frequently associate with RtcR homologs (Supplementary Figure S14, Figure 7) (Himeno et al., 2014, Guyomar et al., 2021).

**Figure 7:**
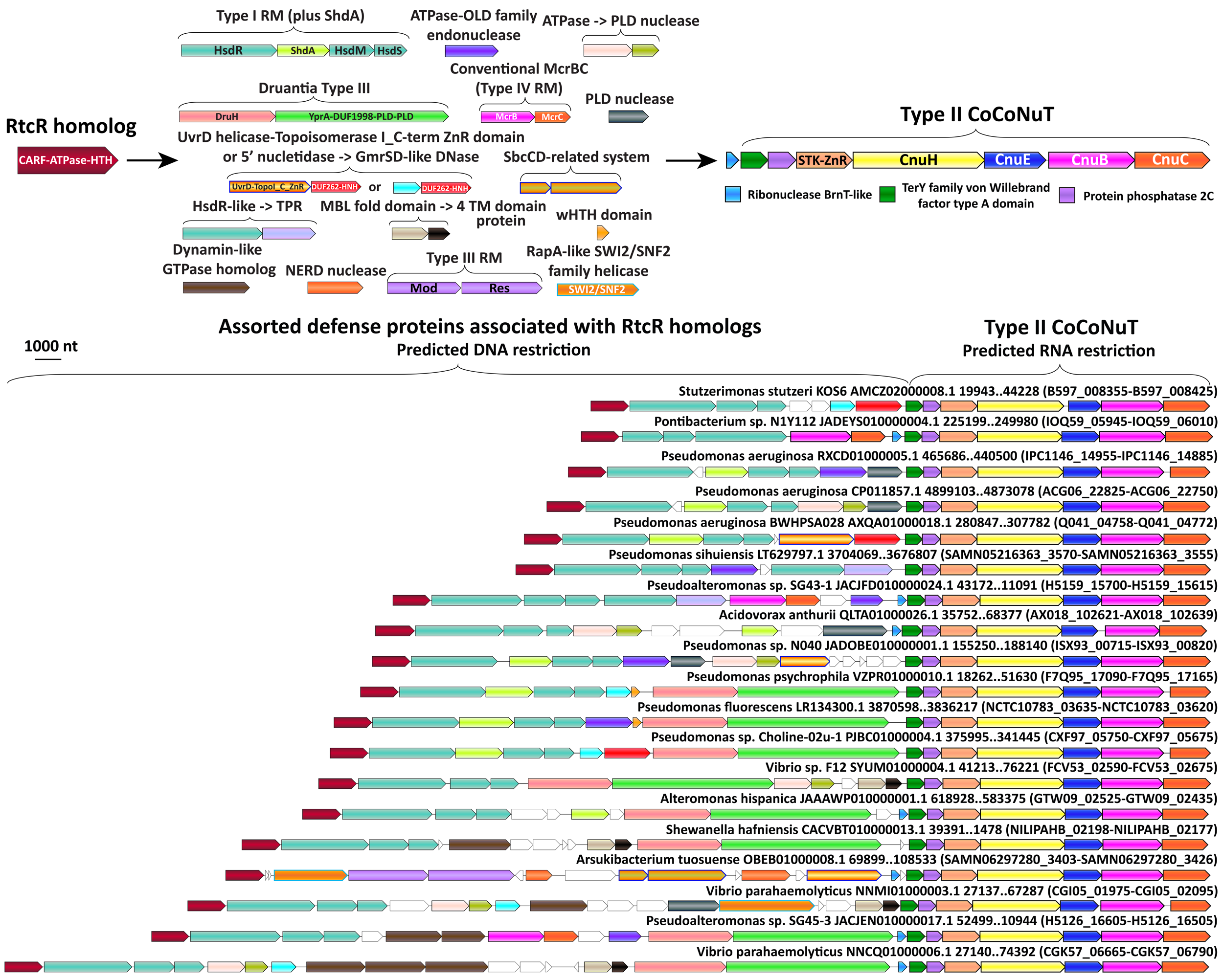
Complex operonic associations of Type II CoCoNuTs. Type II CoCoNuTs are frequently associated with RtcR homologs, and in many cases, ancillary defense genes are located between the RtcR gene and the CoCoNuT, almost always oriented in the same direction in an apparent superoperon. Abbreviations of domains: YprA – YprA-like helicase domain; DUF1998 - DUF1998 is often found in or associated with helicases and contains four conserved, putatively metal ion-binding cysteine residues; PLD – Phospholipase D family nuclease domain; SWI2/SNF2 – SWI2/SNF2-family ATPase; HsdR/M/S – Type I RM system restriction, methylation, and specificity factors; ShdA – Shield system core component ShdA; TPR – Tetratricopeptide repeat protein; MBL fold – Metallo-beta-lactamase fold; 4 TM domain – Protein with 4 predicted transmembrane helices; Mod/Res – Type III RM modification and restriction factors.

There are notable parallels between CoCoNuTs and Type III CRISPR-Cas systems, where the Cas10-Csm-crRNA effector complex binds phage RNA complementary to the spacer of the crRNA, triggering both restriction of phage DNA and indiscriminate cleavage of RNA (McMahon et al., 2020). The target RNA recognition stimulates the production of cyclic oligoadenylate (cOA) signals by Cas10, which activate, via CARF domain binding, PCD effectors, typically HEPN domain-containing proteins, such as Csm6, that function as promiscuous RNases (Pillon et al., 2021, Millman et al., 2020). One of the two outcomes can result from this cascade: the infection can either be eradicated quickly due to the restriction of the virus DNA, which inhibits cOA signaling via the depletion of viral RNA, along with the activity of RING nucleases, thus averting PCD, or the continued presence of viral RNA stimulates cOA signaling until PCD or dormancy occurs, limiting the spread of viruses to neighboring cells in the bacterial population (Makarova et al., 2012, Koonin and Aravind, 2002, Koonin and Krupovic, 2019).

If CoCoNuTs associated with RtcR homologs can induce PCD, a conceptually similar but mechanistically distinct phenomenon might occur. Although many of these CoCoNuTs only contain an appended gene encoding a CARF domain-containing protein at the 5’ end of the predicted operon (Supplementary Figure S14), there are also numerous cases where several types of DNA restriction systems are encoded between the CARF gene and the CoCoNuT (Figure 7). In these cases, nearly all genes are in an apparent operonic organization that can extend upwards of 40 kb (Figure 7). Although internal RtcR-independent promoters likely exist in these large loci, the consistent directionality and close spacing of the genes in these superoperons suggests coordination of expression. The complex organization of the CARF-CoCoNuT genomic regions, and by implication, the corresponding defense mechanisms, might accomplish the same effect as Type III CRISPR-Cas, contriving a no-win situation for the target virus. Under this scenario, the virus is either destroyed by the activity of the DNA restriction systems, which would inhibit signal production (likely RNA fragments with cyclic phosphate ends rather than cOA) and drive down CoCoNuT transcription, thereby preventing PCD, or as the virus replicates, signaling and CoCoNuT transcription would continue until the infection is aborted by PCD or dormancy caused by the degradation of host mRNA via the HEPN RNase(s) of the CoCoNuT.

In many species of *Pseudomonas*, where CoCoNuTs are almost always associated with RtcR and various ancillary factors, Type I RM systems often contain an additional gene that encodes a transmembrane helix and a long coiled-coil fused to an RmuC-like nuclease (Figure 7). These proteins were recently described as ShdA, the core component of the *Pseudomonas-* specific defense system Shield (Macdonald et al., 2022). These can potentially interact with coiled-coils in the CoCoNuTs, perhaps, guiding them to the DNA from which RNA targeted by the CoCoNuTs is being transcribed, or vice versa (Figure 7).

A notable difference between the generally similar, dual DNA and RNA-targeting mechanism of many Type III CRISPR-Cas systems and the proposed mechanism of the CoCoNuTs is that viral RNA recognition by the Cas10-Csm complex, rather than binding of a second messenger to a CARF domain, activates both DNA cleavage activity by the HD domain and production of cOA that triggers non-specific RNA cleavage. In contrast, in the CARF-containing CoCoNuTs, both the DNA and RNA restriction factors appear to be arranged such that binding of a 2’,3’ cyclic phosphate RNA fragment by the CARF domain would initiate the expression of the entire cluster (McMahon et al., 2020). In the case of the CoCoNuTs, signals of infection could promote transcription, first of the DNA restriction systems, and then, the CoCoNuT itself, a predicted RNA restriction system. In these complex configurations of the CoCoNuT genome neighborhoods (Figure 7), the gene order is likely to be important, with Type I RM almost always directly following CARF genes and CoCoNuTs typically coming last, although Druantia Type III sometimes follows the CoCoNuT. As translation in bacteria is co-transcriptional, the products of genes transcribed first would accumulate before those of the genes transcribed last, so that the full, potentially suicidal impact of the CoCoNuT predicted RNA nucleolytic engine would only be felt after the associated DNA restriction systems had ample time to act – and possibly, fail (Irastortza-Olaziregi and Amster-Choder, 2020).

#### Concluding remarks

In recent years, systematic searches for defense systems in prokaryotes, primarily by analysis of defense islands, revealed enormous, previously unsuspected diversity of such systems that function through a remarkable variety of molecular mechanisms. In this work, we uncovered the hidden diversity and striking hierarchical complexity of a distinct class of defense mechanisms, the Type IV (McrBC) restriction systems. We then zeroed in on a single major but previously overlooked branch of the McrBC systems, which we denoted CoCoNuTs for their salient features, namely, the presence of extensive coiled-coil structures and tandem nucleases. Astounding complexity was discovered at this level as well, with 3 distinct types and multiple subtypes of CoCoNuTs that differ by their domain compositions and genomic associations. All CoCoNuTs contain domains capable of interacting with translation systems components, such as the SmpB-like OB-fold, Hsp70 homologs, or YTH domains, along with RNases, such as HEPN, suggesting that at least one of the activities of these systems targets RNA. Most of the CoCoNuTs are likely endowed with DNA-targeting activity as well, either by factors integral to the system, such as the McrC nuclease, or more loosely associated, such as Type I RM and Druantia Type III systems that are encoded in the same predicted superoperon with many CoCoNuTs. Numerous CoCoNuTs are associated with proteins containing CARF domains, suggesting that cyclic (oligo)nucleotides regulate the CoCoNuT activity. Given the presence of the RtcR-like CARF domains, it appears likely that the specific second messengers involved are RNA fragments with cyclic phosphate termini. We hypothesize that the CoCoNuTs, in conjunction with ancillary restriction factors, implement an echeloned defense strategy analogous to that of Type III CRISPR-Cas systems, whereby an immune response eliminating virus DNA and/or RNA is launched first, but then, if it fails, an abortive infection response leading to PCD/dormancy via host RNA cleavage takes over.

## Methods

### Comprehensive identification and phylogenetic and genomic neighborhood analysis of McrB and McrC proteins

The comprehensive search for McrB and McrC proteins was seeded with publicly available multiple sequence alignments COG1401 (McrB GTPase domain), AAA_5 (The branch of AAA+ ATPases containing the McrB GTPase), COG4268 (McrC), PF10117 (McrBC), COG1700 (McrC PD-D/ExK nuclease domain), PF04411 (McrC PD-D/ExK nuclease domain), PF09823 (McrC N-terminal DUF2357). Additional alignments and individual queries were derived from data from our previous work on the EVE domain family (Bell et al., 2020). All alignments were clustered, and each sub-alignment or individual query sequence was used to produce a position-specific scoring matrix (PSSM). These PSSMs were used as PSI-BLAST queries against the non-redundant (nr) NCBI database (*E*-value ≤L10) (Altschul et al., 1997). Although a branch of McrB GTPase homologs has been described in animals, these are highly divergent in function, and no associated McrC homologs have been reported (Iyer et al., 2004). Therefore, we excluded eukaryotic sequences from our analysis to focus on the composition and contextual connections of prokaryotic McrBC systems.

Genome neighborhoods for the hits were generated by downloading their gene sequence, coordinates, and directional information from GenBank, as well as for 10 genes on each side of the hit. Domains in these genes were identified using PSI-BLAST against alignments of domains in the NCBI Conserved Domain Database (CDD) and Pfam (*E*-value 0.001). Some genes were additionally analyzed with HHpred for validation of the BLAST hits or if no hits were obtained. (Gabler et al., 2020). Then, these neighborhoods were filtered for the presence of a COG1401 hit (McrB GTPase domain), or hits to both an McrB “alias” (MoxR, AAA_5, COG4127, DUF4357, Smc, WEMBL, Myosin_tail_1, DUF3578, EVE, Mrr_N, pfam01878) and an McrC “alias” (McrBC, McrC, PF09823, DUF2357, COG1700, PDDEXK_7, RE_LlaJI). These aliases were determined from a preliminary manual investigation of the data using HHpred (Zimmermann et al., 2018a). Several of the McrB aliases are not specific to McrB, and instead are domains commonly fused to McrB GTPase homologs, or are larger families, such as AAA_5, that contain McrB homologs. We found that many *bona fide* McrB homologs, validated by HHpred, produced hits not to COG1401 but rather to these other domains, so we made an effort to retain them.

The McrB candidates identified by this filtering process were clustered to a similarity threshold of 0.5 with MMseqs2 (Hauser et al., 2016), after which the sequences in each cluster were aligned with MUSCLE (Edgar, 2004). Next, profile-to-profile similarity scores between all clusters were calculated with HHsearch (Söding, 2004). Clusters with high similarity, defined as a pairwise score to self-score ratio >L0.1, were aligned to each other with HHalign (Söding et al., 2006). This procedure was performed for a total of 3 iterations. The alignments of each cluster resulting from this protocol, which included some false positive clusters consisting of other members of the AAA_5 family, were analyzed with HHpred to remove the false positives, after which the GTPase domain sequences were extracted manually using HHpred, and the alignments were used as queries for a second round of PSI-BLAST against the nr NCBI database as described above. At this stage, the abundance of the CoCoNuT and CoCoPALM (<u>co</u>iled-<u>co</u>il and <u>p</u>ilus <u>a</u>ssembly linked to <u>M</u>crBC, see above) types of McrB GTPases had become apparent, therefore, results of targeted searches for these subtypes were included in the pool of hits from the second round of PSI-BLAST.

Genome neighborhoods were generated for these hits and filtered for aliases as described above. Further filtering of the data, which did not pass this initial filter, involved relaxing the criteria to include neighborhoods with only one hit to an McrB or McrC alias, but with a gene adjacent to the hit (within 90 nucleotides), oriented in the same direction as the hit, and encoding a protein of sufficient size (> 200 aa for McrB, > 150 aa for McrC) to be the undetected McrB or McrC component. Afterward, we filtered the remaining data to retain genome islands with no McrBC aliases but with PSI-BLAST hits in operonic association with genes of sufficient size, as described above, to be the other McrBC component. These data, which contained many false positives but captured many rare variants, were then clustered and analyzed as described above to remove false positives. Next, an automated procedure was developed to excise the GTPase domain sequences using the manual alignments generated during the first phase of the search as a reference. These GTPase sequences were further analyzed by clustering and HHpred to remove false positives.

Definitive validation by pairing McrB homologs with their respective McrC homologs was also used to corroborate their identification. Occasionally, the McrB and McrC homologs were separated by intervening genes, or the operon order was reversed, and consideration of those possibilities allowed the validation of many additional systems. The pairing process was complicated by and drew our attention to the frequent occurrence of multiple copies of McrBC systems in the same islands that may function cooperatively. Lastly, the rigorously validated set of McrBC pairs, supplemented only with orphans manually annotated as McrBC components using HHpred, were used for our phylogenetics. The final alignments of GTPase and DUF2357 domains were produced using the iterative alignment procedure described above for 10 iterations. Approximately-maximum-likelihood trees were built with the FastTree program (Price et al., 2010) from representative sequences following clustering to a 0.9 similarity threshold with MMseqs2.

### Domain detection and annotation

Protein domains in McrBC homologs and in proteins encoded by neighboring genes were initially identified using the method described above, the first pass using PSI-BLAST against alignments of domains in the NCBI Conserved Domain Database (CDD) and Pfam (*E*-value 0.001). In many cases where no domains could be confidently detected with this method, or for validation of the hits from the first pass, HHpred was used for more sensitive analysis (Zimmermann et al., 2018b). The CoCoNuT system components were subjected to additional scrutiny using the coiled-coil detection and visualization tool Waggawagga, which employs a several algorithms for coiled-coil prediction, including Marcoil, Multicoil2, Ncoils, and Paircoil2 (Simm et al., 2015, Delorenzi and Speed, 2002, Trigg et al., 2011, Lupas et al., 1991, McDonnell et al., 2006). These predictions varied in their strength, with the long coiled-coils detected in CnuA and CnuB homologs having the highest likelihood (usually the maximum P-score of 100 with Marcoil and Multicoil2) and being recognized by the most of the applied tools (BLAST, HHpred, and multiple algorithms used by Waggawagga). The shorter coiled-coils in CnuC, CnuD, and CnuH homologs were less strongly, but nevertheless confidently predicted, usually being detected by HHpred and by at least one but typically, more than one, coiled-coil prediction tool. The analysis was performed on both representative individual sequences and consensus sequences. The potential coiled-coils in CnuE homologs were often only found by Ncoils and were near the limit of detection, but these regions were also reported as coiled-coils in another study (Anantharaman et al., 2012). Given the context of extensive, high-probability coiled-coils in other components of the CoCoNuT systems with which they might interact, we chose to report these CnuE regions as coiled-coils, despite the comparative weakness of these predictions. In the AF2 multimer model of CnuHE, one of these potential coiled-coils is positioned near the coiled-coils detected in CnuH, suggesting they may facilitate interaction between these two factors.

### Preliminary phylogenetic analysis of CoCoNuT CnuH helicases

A comprehensive search and phylogenetic analysis of this family was beyond the scope of this work, but to determine the relationships between the CoCoNuTs and the rest of the UPF1-like helicases, we used the following procedure. We retrieved the best 2000 hits in each of two searches with UPF1 and CoCoNuT helicases as queries against both a database containing predicted proteins from 24,757 completely sequenced prokaryotic genomes downloaded from the NCBI GenBank in November 2021 and a database containing 72 representative eukaryotic genomes that were downloaded from the NCBI GenBank in June 2020. Next, we combined all proteins in four searches, made a nonredundant set, and annotated them using CDD profiles, as described above. Then, we aligned them with MUSCLE v5 (Robert, 2022), constructed an approximately-maximum-likelihood tree with FastTree, and mapped the annotations onto the tree. Genome neighborhoods were generated for these hits, as described above.

### Structural modeling with AlphaFold2 and searches for related folds

Protein structures were predicted using AlphaFold2 (AF2) v2.2.0 with local installations of complete databases required for AF2 (Jumper et al., 2021). Only protein models with average predicted local distance difference test (pLDDT) scores > 80 were retained for further analysis (Mariani et al., 2013). Many of these models were used as queries to search for structurally similar proteins using DALI v5 against the Protein Data Bank (PDB) and using FoldSeek against the AlphaFold/UniProt50 v4, AlphaFold/Swiss-Prot v4, AlphaFold/Proteome v4, and PDB100 2201222 databases (Holm, 2020, van Kempen et al., 2023). Structure visualizations and comparisons were performed with ChimeraX (Pettersen et al., 2021) and the RCSB PDB website (Berman et al., 2000).

## Supporting information

Suppelmentary table 1

## Data availability

All data generated and analyzed in this study are included in the manuscript and supporting files. The CoCoNuT systems are documented in detail in the Supplementary Data. The AlphaFold2 structures generated and presented in the figures are available at modelarchive.org with accessions listed in Supplementary Table S2.

## Conflict of interest

The authors declare no conflict of interest.

## Author contributions

R.T.B and E.V.K. initiated the project; R.T.B., Y.I.W., and E.V.K. designed research; R.T.B., H.S., and K.S.M. analyzed data; R.T.B and E.V.K. wrote the manuscript that was edited and approved by all authors. The authors declare no competing interests.

## Acknowledgments

The authors thank Becky Xu Hua Fu (University of California, San Francisco) for correspondence that led to her contribution of the name CoCoNuT, Andrew Z. Fire and Usman Enam (Stanford University) for critical reading of the manuscript and insightful comments, Joseph Bondy-Denomy (University of California, San Francisco) for bringing the Shield factor ShdA to our attention, and Koonin group members for helpful discussions. The authors’ research is supported by the Intramural Research Program of the National Institutes of Health (National Library of Medicine).

## Notes

### Competing Interest Statement

The authors have declared no competing interest.

### Summary of Updates

In this version, the Methods have been amended for clarity and completeness, the Results have been edited for style, and the figures have been improved.

