## Supplementary figures and text for "CoCoNuTs: A diverse subclass of Type IV restriction systems predicted to target RNA"

**Table of Contents**


| Page | Title |
| --- | --- |
| 1 | **Supplementary Figure S1: Phylogenetic tree of the McrC DUF2357 domain** |
| 2 | **Supplementary Figure S2: Phylogeny of McrB family GTPases containing the NxD motif** |
| 3 | **Supplementary Figure S3: Domain composition and AlphaFold2 structural predictions of components of the Type I CoCoNuT systems colored by pLDDT** |
| 4 | **Supplementary Figure S4: Pseudo-Type I-B CoCoNuT genomic context in *Bacillus*** |
| 4 | **Supplementary Note S1: *Functional prediction for Pseudo-Type I-B CoCoNuTs*** |
| 5 | **Supplementary Figure S5: Comparisons of Type II and Type III-A CoCoNuT N-terminal SPB domains, SmpB, and prokaryotic HIRAN domains** |
| 6 | **Supplementary Figure S6: Domain composition and AlphaFold2 structural predictions for core protein components of Type II and III-A CoCoNuT systems colored by pLDDT** |
| 7 | **Supplementary Figure S7: AlphaFold2 prediction of Type I-A CoCoNuT CnuB GTPase hexamer and CnuC monomer complex** |
| 8 | **Supplementary Figure S8: AlphaFold2 prediction of Type I-B CoCoNuT CnuB hexamer and CnuC monomer complex** |
| 9 | **Supplementary Figure S9: AlphaFold2 prediction of Type I-C CoCoNuT CnuB GTPase hexamer and CnuC monomer complex** |
| 10 | **Supplementary Figure S10: AlphaFold2 prediction of Type II CoCoNuT CnuB GTPase hexamer and CnuC monomer complex** |
| 11 | **Supplementary Figure S11: AlphaFold2 prediction of Type II CoCoNuT CnuH helicase and CnuE effector complex** |
| 12 | **Supplementary Figure S12: AlphaFold2 prediction of Type III-A CoCoNuT CnuB GTPase hexamer and CnuC monomer complex** |
| 13 | **Supplementary Figure S13: Alignment of Hsp70-like NBDs from CoCoNuTs and related McrB homolog with *E. coli* Hsp70 (DnaK) and mammalian Hsp70 cognate protein *H. sapiens* HSC70** |
| 14 | **Supplementary Figure S14: Type II CoCoNuTs are associated with RtcR homologs in a variety of species** |
| 15-17 | **Supplementary Note S2: *Extension of Type III-A CoCoNuT systems with ATPases and virulence factors*** |
| 15 | **Supplementary Figure S15: Compound Type III-A CoCoNuT operons with 5' extensions** |
| 18-19 | **Supplementary References** |

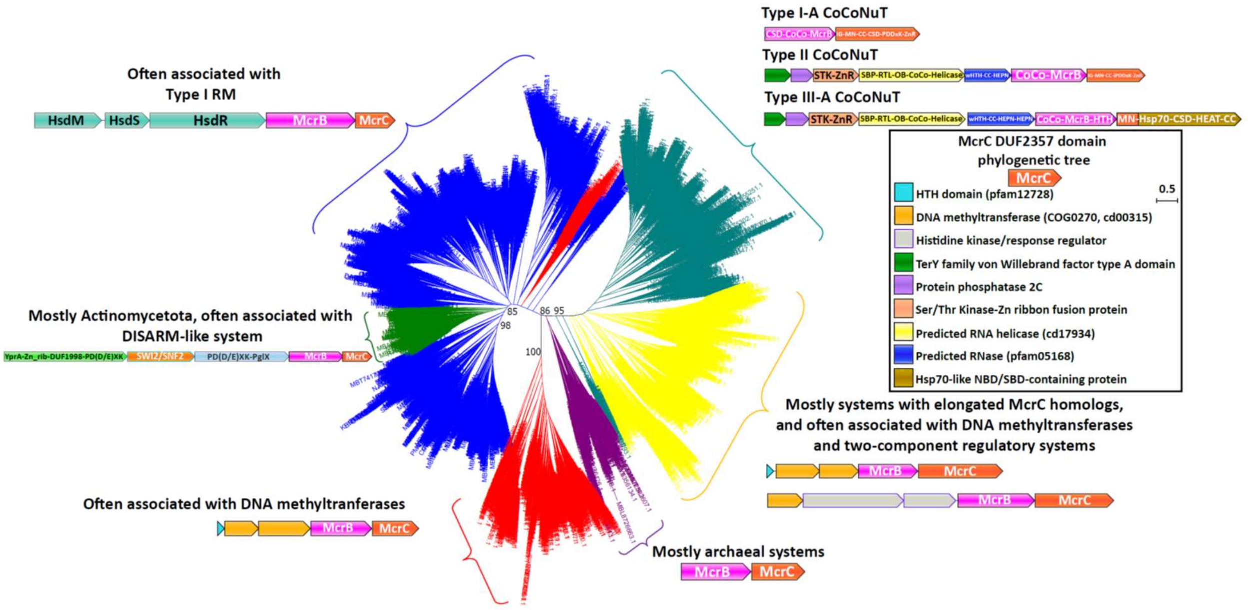


**Supplementary Figure S1: Phylogenetic tree of the McrC DUF2357 domain**

The phylogenetic tree of the McrC N-terminal DUF2357 domains is generally topologically concordant with the McrB GTPase tree. Each of the differently colored groups is characterized by distinct conserved genomic associations that are abundant within but not completely confined to the respective groups. This tree was built from the representatives of 90% identity clusters of all validated homologs. Abbreviations of domains: McrB – McrB GTPase domain; CoCo/CC – coiled-coil; MN – McrC N-terminal domain (DUF2357); CSD – cold shock domain; IG – Immunoglobulin-like beta-sandwich domain; ZnR – zinc ribbon domain; SPB – SmpB-like domain; RTL – RNase toxin-like domain; HEPN – HEPN family nuclease domain; OB – OB-fold domain; iPD-DxK – inactivated PD-DxK fold; Hsp70 – Hsp70-like NBD/SBD; HEAT – HEAT-like helical repeats; YprA – YprA-like helicase domain; DUF1998 - DUF1998 is often found in or associated with helicases and contains four conserved, putatively metal ion-binding cysteine residues; SWI2/SNF2 – SWI2/SNF2-family ATPase; PglX – PglX-like DNA methyltransferase; HsdR/M/S – Type I RM system restriction, methylation, and specificity factors.

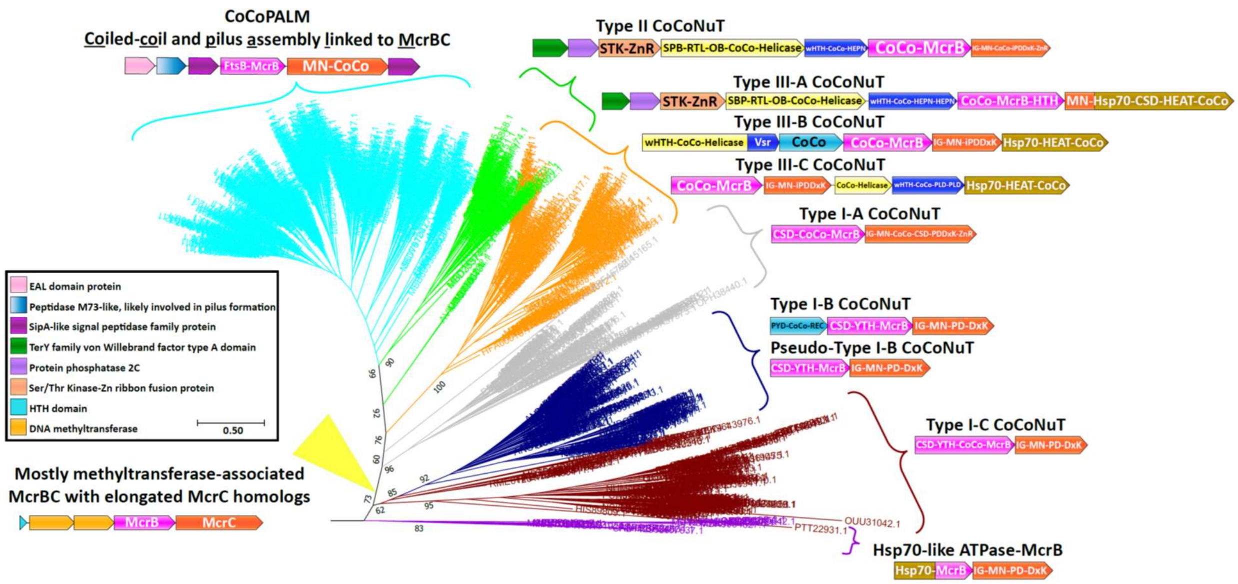


**Supplementary Figure S2: Phylogeny of McrB family GTPases containing the NxD motif**

The phylogenetic tree of McrB GTPases with an NxD variant of the signature motif contains the CoCoNuTs, CoCoPALMs, and systems associated with methyltransferases with additional domains fused to the McrC homologs. Each of the differently colored branches is characterized by distinct conserved genomic associations and domain compositions, which we used to define 3 CoCoNuT types and 7 subtypes. These types are not generally found in other branches of the tree, except some CoCoPALMs, which can be found in the Type II CoCoNuT branch. This tree was built from the representatives of 90% identity clusters of all validated homologs. Abbreviations of domains: McrB – McrB GTPase domain; CoCo/CC – coiled-coil; MN – McrC N-terminal domain (DUF2357); CSD – cold shock domain; YTH – YTH-like domain; IG – Immunoglobulin (IG)-like beta-sandwich domain; Hsp70 – Hsp70-like NBD/SBD; HEAT – HEAT-like helical repeats; ZnR – zinc ribbon domain; PYD – pyrin/CARD-like domain; SPB – SmpB-like domain; RTL – RNase toxin-like domain; HEPN – HEPN family nuclease domain; OB – OB-fold domain; wHTH – winged helix-turn-helix (HTH) domain; iPD-DxK – inactivated PD-DxK fold; FtsB – FtsB-like TM helix and CoCo; REC – Phosphoacceptor receiver-like domain; PLD – Phospholipase D-like nuclease domain; Vsr – very-short-patch-repair nuclease-like domain.

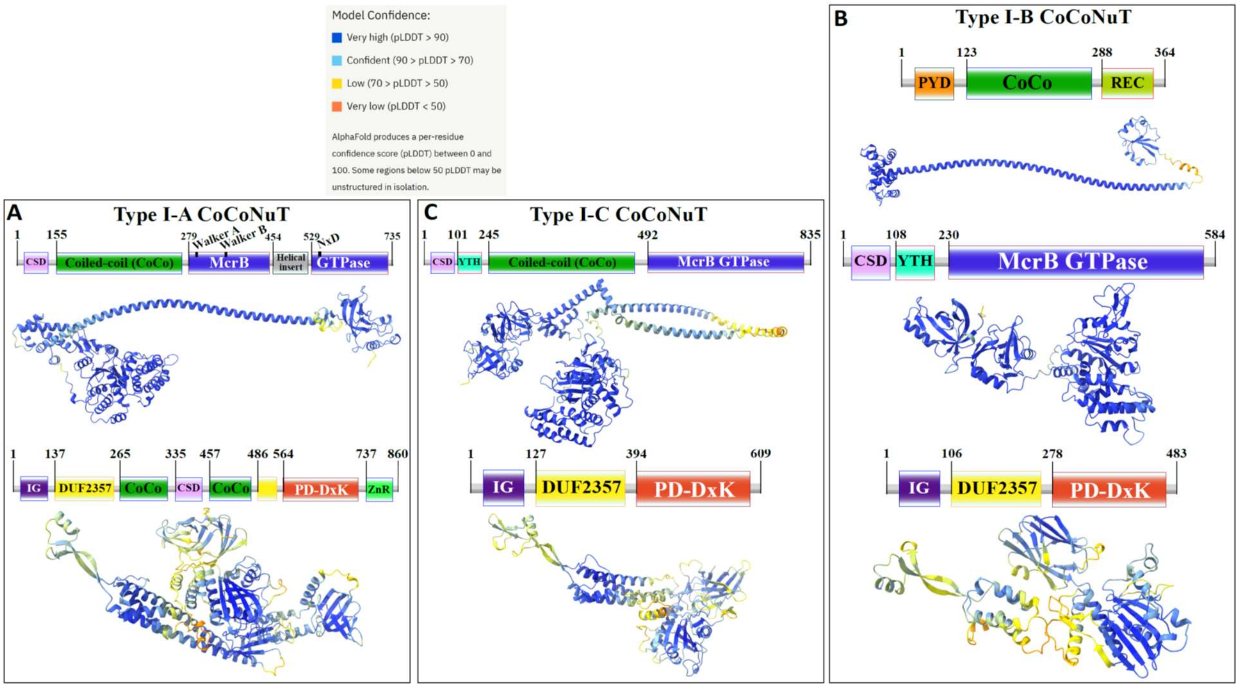


**Supplementary Figure S3: Domain composition and AlphaFold2 structural predictions of components of the Type I CoCoNuT systems colored by pLDDT**

A-C) High quality (Average pLDDT > 80), representative AlphaFold2 structural predictions for proteins in A) Type I-A CoCoNuT systems (CnuB and CnuC, from top to bottom), B) Type I-B (CnuA, CnuB, and CnuC, from top to bottom), and C) Type I-C CoCoNuT systems (CnuB and CnuC, from top to bottom). Models were generated from representative sequences with the following GenBank accessions (see Supplementary Data for sequences and locus tags): ROR86958.1 (Type I-A CnuB), APL73566.1 (Type I-A CnuC), TKH01449.1 (Type I-B CnuA), GED20858.1 (Type I-B CnuB), GED20857.1 (Type I-B CnuC), GFD85286.1 (Type I-C CnuB), MBV0932851.1 (Type I-C CnuC). Abbreviations of domains: CSD – cold shock domain; YTH – YTH-like domain; CoCo – coiled-coil; IG – Immunoglobulin (IG)-like beta-sandwich domain; ZnR – zinc ribbon domain; PYD – pyrin/CARD-like domain; REC – Phosphoacceptor receiver-like domain. These structures were visualized with ChimeraX [1].

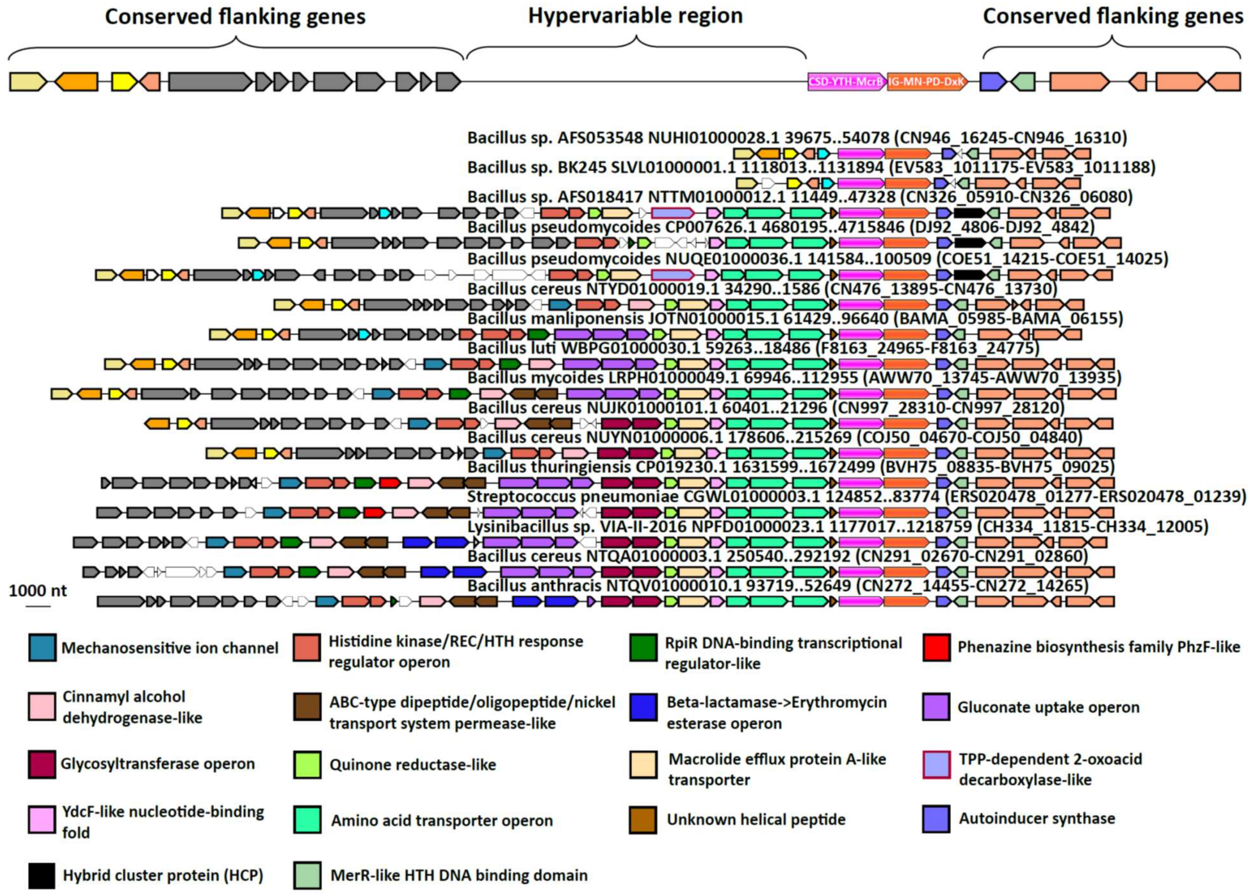


**Supplementary Figure S4: Pseudo-Type I-B CoCoNuT genomic context in *Bacillus***

Pseudo-Type I-B CoCoNuTs in *Bacillus* are associated with various factors with potential involvement in overcrowding-induced stress. Abbreviations of domains: McrB – McrB GTPase domain; MN – McrC N-terminal domain (DUF2357); CSD – cold shock domain; YTH – YTH-like domain; IG – Immunoglobulin (IG)-like beta-sandwich domain.

### Supplementary Note S1

*Functional prediction for Pseudo-Type I-B CoCoNuTs*

We hypothesize that Pseudo-Type I-B CoCoNuTs play a role in overcrowding-induced stress responses. We inferred this functional prediction from the fact that the islands in which it occurs typically also encode a quorum-sensing hormone synthase, a mechanosensitive ion channel, various transporters, antibiotic resistance and synthesis factors, and cell wall-related proteins. In many cases, we detected Type I-B CoCoNuT homologs in the extended neighborhoods of the Pseudo-Type I-B CoCoNuTs, although only rarely in the immediate vicinity. Thus, Pseudo-Type I-B CoCoNuTs might be derived duplicates of Type I-B CoCoNuTs that acquired a specialized but likely related functionality, perhaps still using the coiled-coil protein encoded by the Type I-B CoCoNuT.

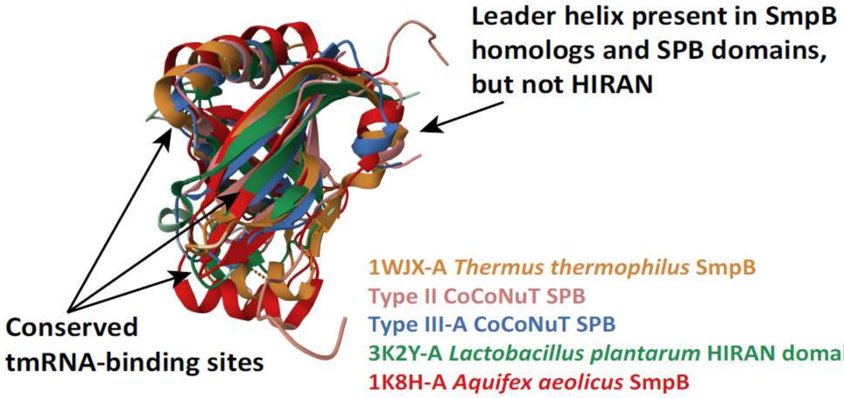


| | RMSD vs. 1WJX-A | TM-score vs. 1WJX-A | Identity vs. 1WJX-A |
|---|---|---|---|
| 1WJX-A *Thermus thermophilus* SmpB | | | |
| Type II CoCoNuT SPB | 3.13 | 0.54 | 2% |
| Type III-A CoCoNuT SPB | 3.03 | 0.55 | 6% |
| 3K2Y-A *Lactobacillus plantarum* HIRAN domain | 3.35 | 0.53 | 10% |
| 1K8H-A *Aquifex aeolicus* SmpB | 2.92 | 0.62 | 62% |

**Supplementary Figure S5: Comparisons of Type II and Type III-A CoCoNuT N-terminal SPB domains, SmpB, and prokaryotic HIRAN domains**

High quality (Average pLDDT > 80) AlphaFold2 representative structural predictions for the N-terminal SPB domains in Type II and III-A helicases and experimentally solved structures for SmpB and a prokaryotic HIRAN domain. Models were generated from representative sequences with the following GenBank accessions (see Supplementary Data for sequences and locus tags): AMO81399.1 (Type II SPB), PJX13386.1 (Type III-A SPB). Structures were visualized and compared using the pairwise alignment tool on the RCSB PDB website [2].

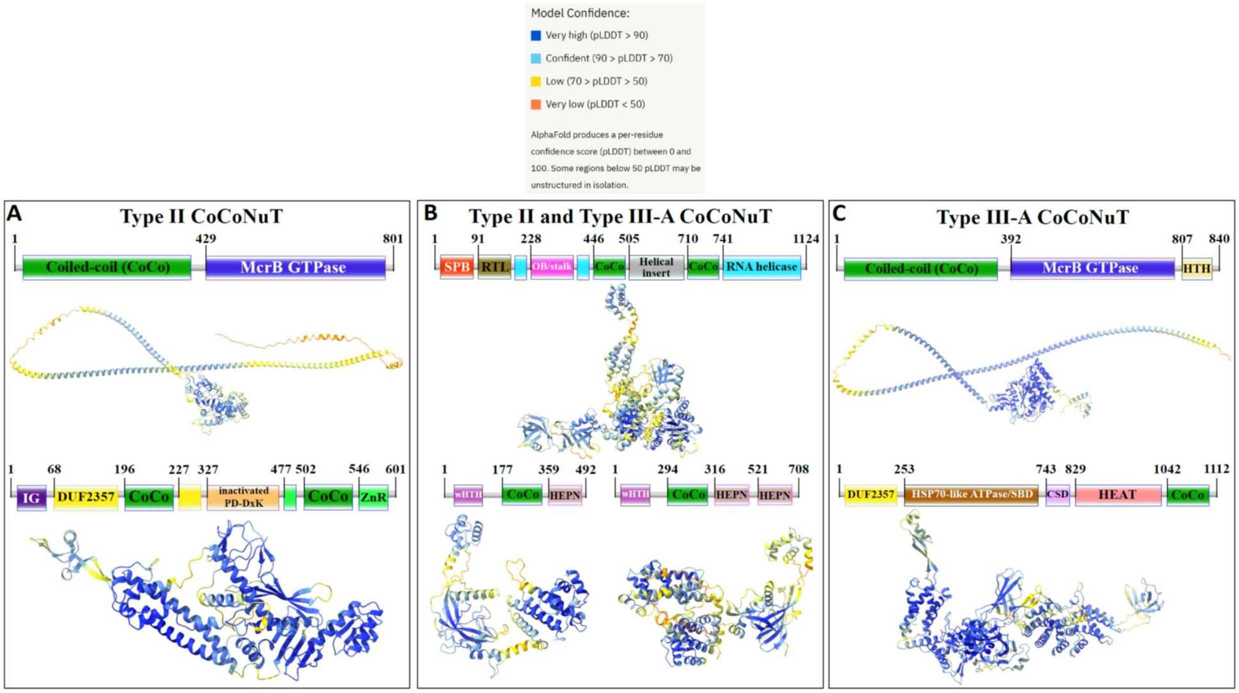


**Supplementary Figure S6: Domain composition and AlphaFold2 structural predictions for core protein components of Type II and III-A CoCoNuT systems colored by pLDDT**

A-C) High quality (Average pLDDT > 80), representative AlphaFold2 structural predictions for proteins in A) Type II CoCoNuT systems (CnuB and CnuC, from top to bottom), B) Type II and III-A CoCoNuT systems (CnuH at the top, Type II CnuE on the bottom left, Type III-A CnuE on the bottom right), and C) Type III-A CoCoNuT systems (CnuB and CnuC, from top to bottom). Models were generated from representative sequences with the following GenBank accessions (see Supplementary Data for sequences and locus tags): AMO81401.1 (Type II CnuB), AVE71177.1 (Type II CnuC), AMO81399.1 (Type II and III-A CoCoNuT CnuH), AVE71179.1 (Type II CnuE), ATV59464.1 (Type III-A CnuE), PNG83940.1 (Type III-A CnuB), NMY00740.1 (Type III-A CnuC). Abbreviations of domains: CSD – cold shock domain; CoCo – coiled-coil; IG – Immunoglobulin (IG)-like beta-sandwich domain; ZnR – zinc ribbon domain; SPB – SmpB-like domain; RTL – RNase toxin-like domain; HEPN – HEPN family nuclease domain; HTH – helix-turn-helix domain; OB/stalk – OB-fold domain attached to a helical stalk-like extension of ATPase; HEAT – HEAT-like helical repeats. These structures were visualized with ChimeraX [1].

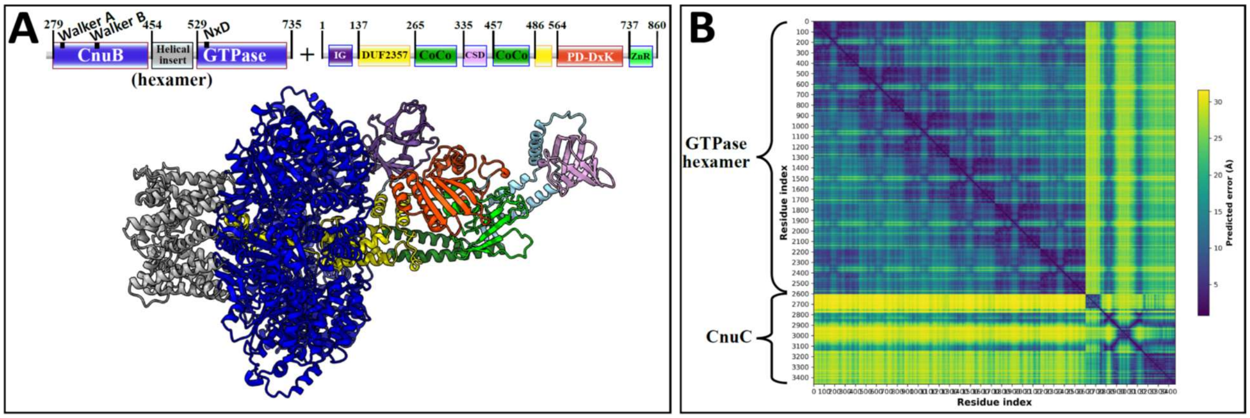


**Supplementary Figure S7: AlphaFold2 prediction of Type I-A CoCoNuT CnuB GTPase hexamer and CnuC monomer complex**

A) High quality (Average pLDDT = 82.5, ipTM+pTM = 0.7392) AlphaFold2 multimer structural prediction for the CnuB GTPase hexamer (without the N-terminal domains) and CnuC monomer complex in a Type I-A CoCoNuT system. B) Predicted aligned error (PAE) plot for the predicted complex. The model was generated from representative sequences with the following GenBank accessions (see Supplementary Data for sequences and locus tags): APL73567.1 (Type I-A CoCoNuT CnuB) and APL73566.1 (Type I-A CoCoNuT CnuC). Abbreviations of domains: CSD – cold shock domain; CoCo – coiled-coil; IG – Immunoglobulin (IG)-like beta-sandwich domain; ZnR – zinc ribbon domain. These structures were visualized with ChimeraX [1].

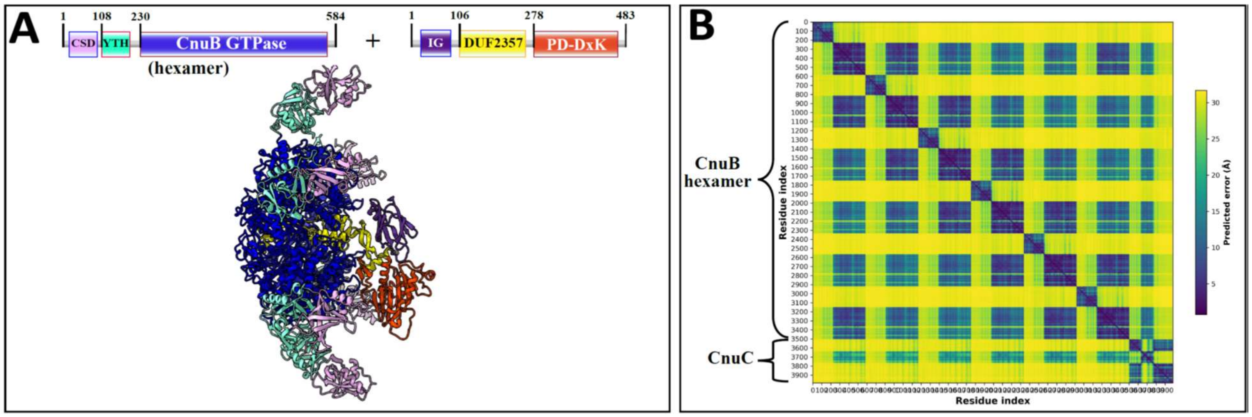


**Supplementary Figure S8: AlphaFold2 prediction of Type I-B CoCoNuT CnuB hexamer and CnuC monomer complex**

A) High quality (Average pLDDT = 77.3, ipTM+pTM = 0.6308) AlphaFold2 multimer structural prediction for the full-length CnuB hexamer and CnuC monomer complex in a Type I-B CoCoNuT system. B) Predicted aligned error (PAE) plot for the predicted complex. The model was generated from representative sequences with the following GenBank accessions (see Supplementary Data for sequences and locus tags): GED20858.1 (Type I-B CoCoNuT CnuB) and GED20857.1 (Type I-B CoCoNuT CnuC). Abbreviations of domains: CSD – cold shock domain; YTH – YTH-like domain; CoCo – coiled-coil; IG – Immunoglobulin (IG)-like beta-sandwich domain. These structures were visualized with ChimeraX [1].

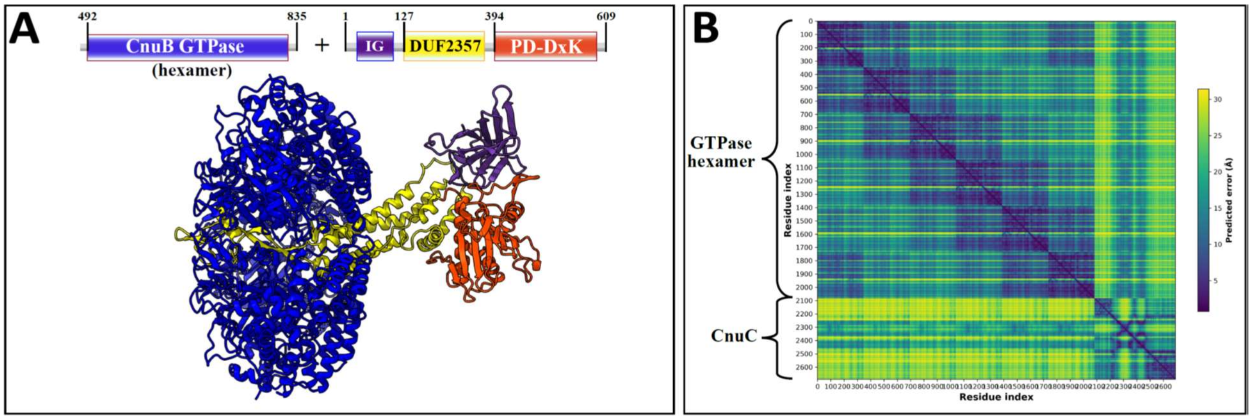


**Supplementary Figure S9: AlphaFold2 prediction of Type I-C CoCoNuT**

**CnuB GTPase hexamer and CnuC monomer complex**

A) High quality (Average pLDDT = 80.0, ipTM+pTM = 0.7271) AlphaFold2 multimer structural prediction for the CnuB GTPase hexamer (without the N-terminal domains) and CnuC monomer complex in a Type I-C CoCoNuT system. B) Predicted aligned error (PAE) plot for the predicted complex. The model was generated from representative sequences with the following GenBank accessions (see Supplementary Data for sequences and locus tags): MBV0932852.1 (Type I-C CoCoNuT CnuB) and MBV0932851.1 (Type I-C CoCoNuT CnuC). Abbreviations of domains: IG – Immunoglobulin (IG)-like beta-sandwich domain. These structures were visualized with ChimeraX [1].

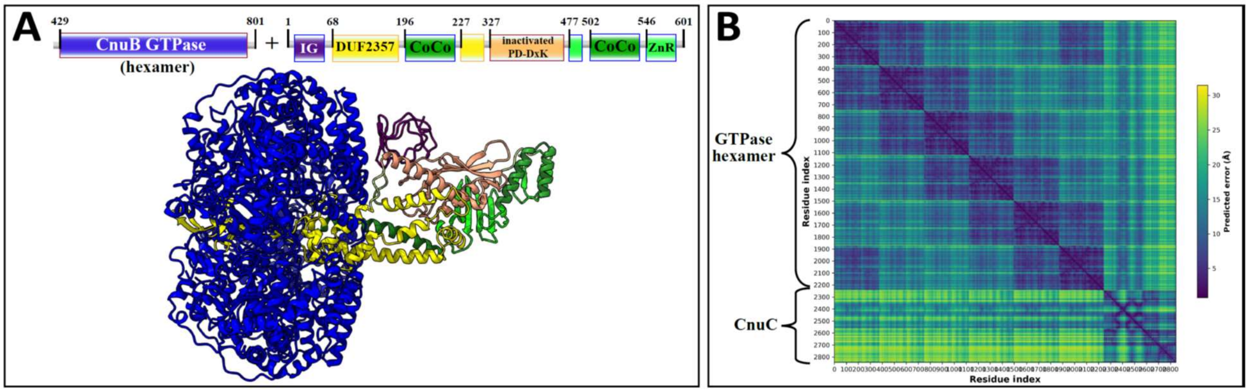


**Supplementary Figure S10: AlphaFold2 prediction of Type II CoCoNuT**

**CnuB GTPase hexamer and CnuC monomer complex**

A) High quality (Average pLDDT = 85.7, ipTM+pTM = 0.7933) AlphaFold2 multimer structural prediction for the CnuB GTPase hexamer (without the N-terminal coiled-coil domain) and CnuC monomer complex in a Type II CoCoNuT system. B) Predicted aligned error (PAE) plot for the predicted complex. The model was generated from representative sequences with the following GenBank accessions (see Supplementary Data for sequences and locus tags): MBV0932852.1 (Type II CoCoNuT CnuB) and MBV0932851.1 (Type II CoCoNuT CnuC). Abbreviations of domains: CoCo – coiled-coil; IG – Immunoglobulin (IG)-like beta-sandwich domain; ZnR – zinc ribbon domain. These structures were visualized with ChimeraX [1].

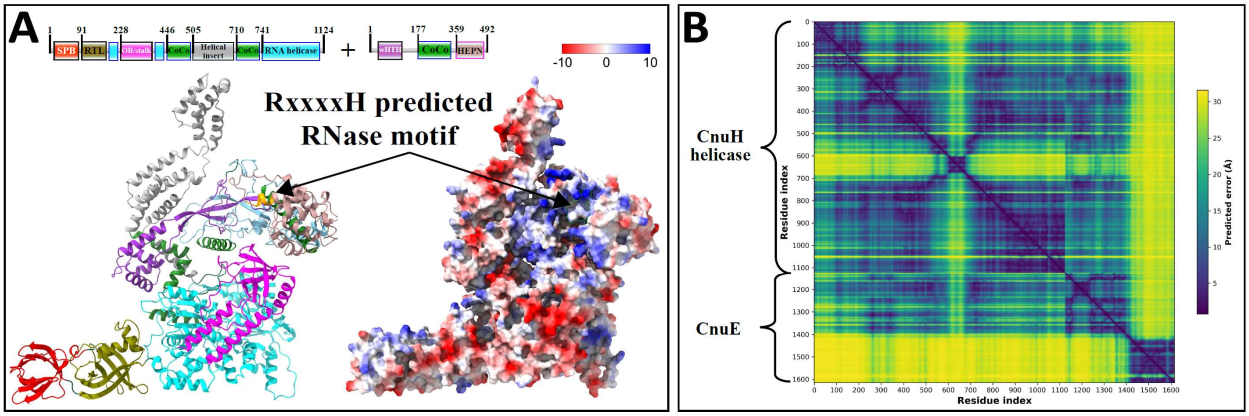


**Supplementary Figure S11: AlphaFold2 prediction of Type II CoCoNuT**

**CnuH helicase and CnuE effector complex**

A) High quality (Average pLDDT = 80.5, ipTM+pTM = 0.7108) AlphaFold2 multimer structural prediction for the CnuH helicase and CnuE wHTH-HEPN effector in a Type II CoCoNuT system. B) Predicted aligned error (PAE) plot for the predicted complex. The helical insert found in the helicase and HEPN domain in the effector show a higher degree of error, but as we predicted them to be novel RNA-binding domains, this is not unexpected. The predicted surface charge distribution shows both domains forming, in conjunction with the OB/stalk domain, a positively charged (blue) groove, with the RxxxxH predicted RNase motif (orange) at its center, that might bind and cleave RNA. The model was generated from representative sequences with the following GenBank accessions (see Supplementary Data for sequences and locus tags): AMO81399.1 (Type II CoCoNuT helicase) and AMO81400.1 (Type II wHTH-CoCo-HEPN protein). Abbreviations of domains: CoCo – coiled-coil; SPB – SmpB-like domain; RTL – RNase toxin-like domain; OB/stalk – OB-fold domain attached to a helical stalk-like extension of ATPase; wHTH – winged helix-turn-helix (HTH) domain; HEPN – HEPN family nuclease domain. These structures were visualized with ChimeraX [1].

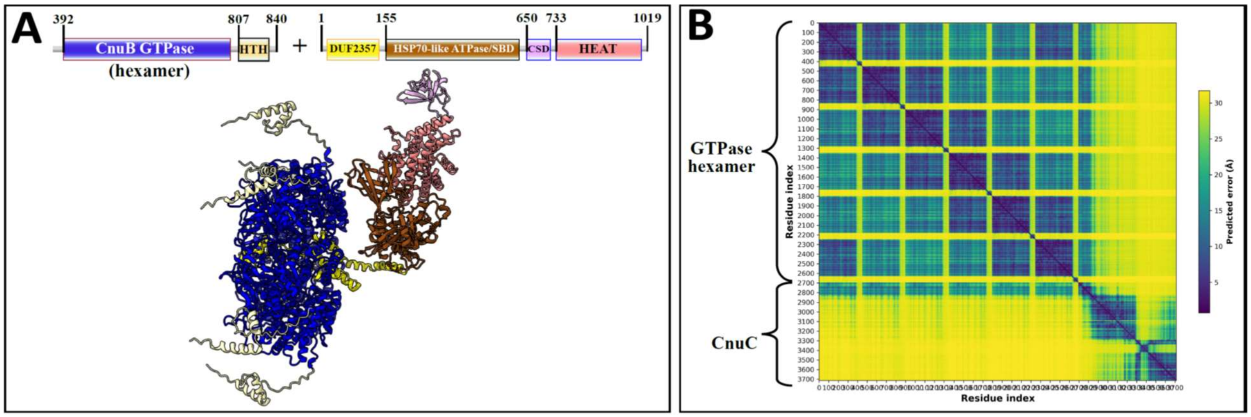


**Supplementary Figure S12: AlphaFold2 prediction of Type III-A CoCoNuT CnuB GTPase hexamer and CnuC monomer complex**

A) High quality (Average pLDDT = 76.6, ipTM+pTM = 0.6472) AlphaFold2 multimer structural prediction for the CnuB GTPase hexamer (without the N-terminal coiled-coil domain) and CnuC monomer complex in a Type III-A CoCoNuT system. B) Predicted aligned error (PAE) plot for the predicted complex. The model was generated from representative sequences with the following GenBank accessions (see Supplementary Data for sequences and locus tags): PNG83940.1 (Type III-A CoCoNuT CnuB) and PNG83939.1 (Type III-A CoCoNuT CnuC). Abbreviations of domains: HTH – helix-turn-helix domain; CSD – cold shock domain; CoCo – coiled-coil; HEAT – HEAT-like helical repeats. These structures were visualized with ChimeraX [1].

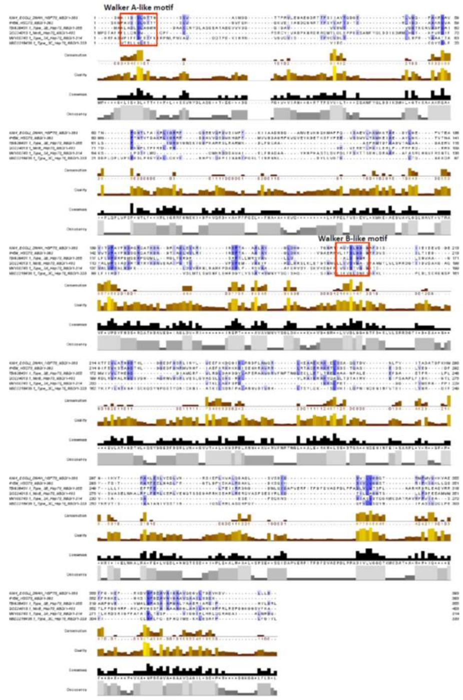


**Supplementary Figure S13: Alignment of Hsp70-like NBDs from CoCoNuTs and related McrB homolog with *E. coli* Hsp70 (DnaK) and mammalian Hsp70 cognate protein *H. sapiens* HSC70**

Alignment of representatives of Hsp70 NBD homologs found in Type III CoCoNuT systems and fused to a closely related NxD motif McrB GTPase with characterized bacterial and mammalian Hsp70 NBD homologs. The alignment was initially performed with PROMALS3D [3] and then adjusted manually. The alignment was displayed with Jalview [4]. The blue shading corresponds to sequence conservation, and the red boxes indicate the Walker A-like and Walker B-like motifs. These motifs are identified on the CDD website in the entry for Hsp70_NBD (cd10170) [5].

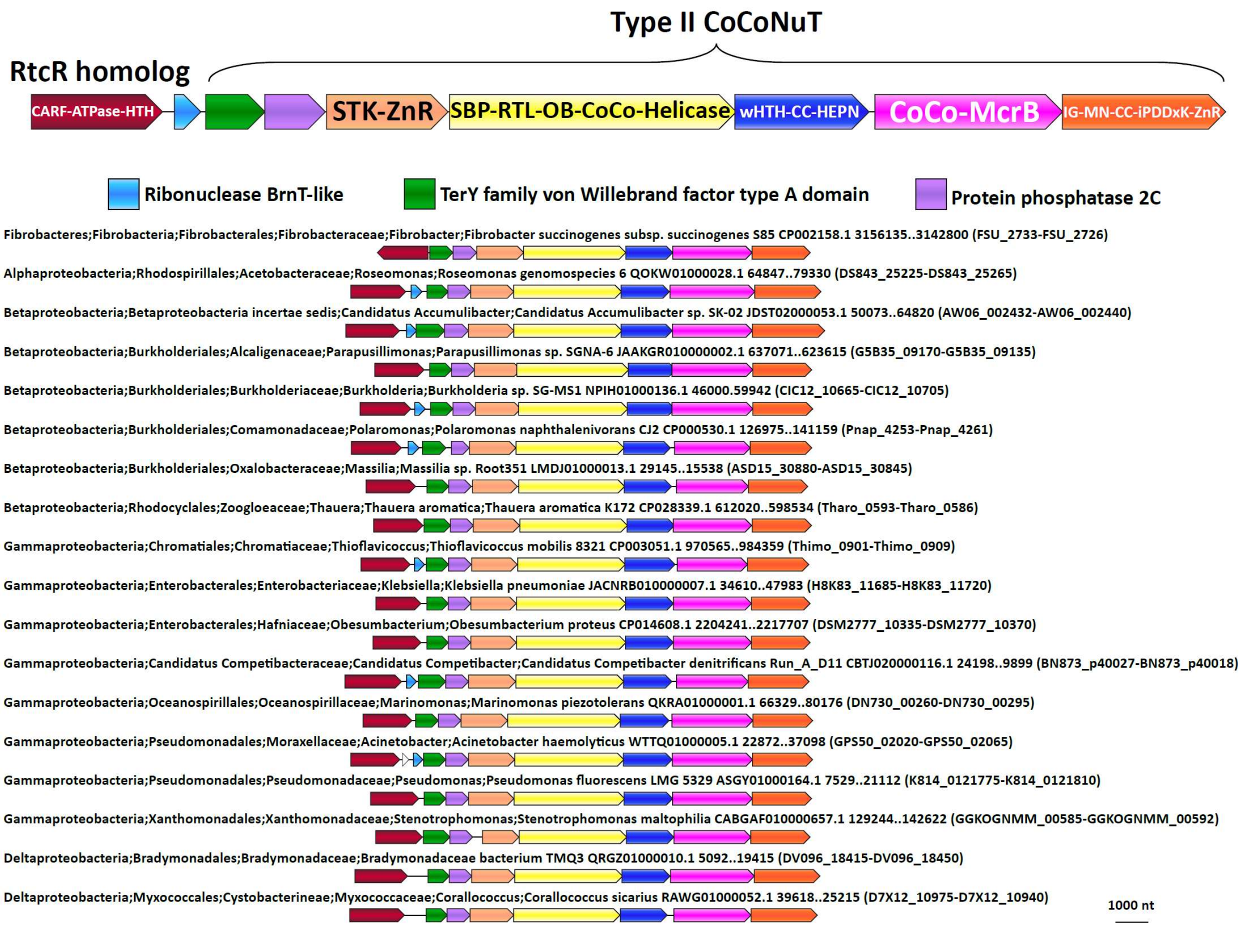


**Supplementary Figure S14: Type II CoCoNuTs are associated with RtcR homologs in a variety of species**

About 30% of Type II CoCoNuT systems detected in this study are associated with RtcR homologs. This contextual connection is conserved in many species of Pseudomonadota.

### Supplementary Note S2

*Extension of Type III-A CoCoNuT systems with ATPases and virulence factors*

We observed even greater levels of complexity in Type III-A CoCoNuTs, which might ultimately beget another level of classification, where the TerY-related VWA domains were duplicated. In each of the 3 subtypes of Type III-A, a different ATPase was inserted between these VWA domains into the predicted operon. In these systems, the $Mg^{2+}$-coordinating MIDAS motif (DxSxS…T...D) is perfectly conserved in the VWA domain encoded at the 5' end of the operon, which resembles the domains found in systems with only one VWA domain, whereas the internal VWA domain lacks the middle threonine and is slightly shorter (Supplementary Figure S15, Supplementary Table S1) [6]. In two of these Type III-A subtypes, a CARF domain-containing protein is encoded, with two distinct CARF domains encompassing different RING nuclease motifs. These CARF domains are fused at the C-terminus to a D-ExK nuclease domain, an architecture suggestive of a PCD/dormancy-eliciting antiphage effector (see main text for a detailed discussion of these and other CARF domains associated with CoCoNuTs) (Supplementary Table S1) [7, 8]. Indeed, homologs of this protein, Can1 and Can2, have been characterized as CRISPR ancillary nucleases [9, 10], and Can2 has been reported to cleave both DNA and RNA [10]. In these two CoCoNuT varieties, the CnuC/McrC homologs typically contain 2 CSDs, whereas in the third type, where the CARF proteins are not encoded, there is a single CSD (Supplementary Figure S15, Supplementary Table S1).

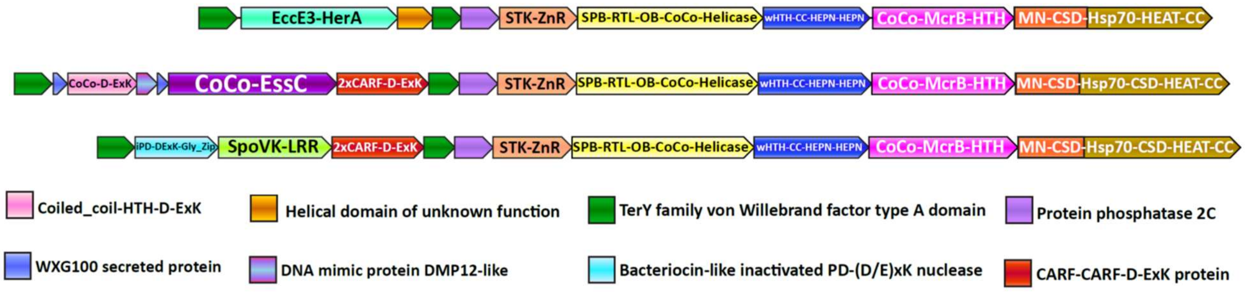


**Supplementary Figure S15: Compound Type III-A CoCoNuT operons with 5' extensions**

Additional genes that may be present in Type III-A CoCoNuT operons. Abbreviations of domains: McrB – McrB GTPase domain; CoCo/CC – coiled-coil; STK – Serine/threonine kinase; 2xCARF – 2 CARF domains; D-ExK – D-ExK nuclease motif; MN – McrC N-terminal domain (DUF2357); CSD – cold shock domain; ZnR – zinc ribbon domain; SPB – SmpB-like domain; RTL – RNase toxin-like domain; OB – OB-fold domain attached to helical stalk-like extension of ATPase; HEPN – HEPN family nuclease domain; Hsp70 – Hsp70-like NBD/SBD; HEAT – HEAT-like helical repeats; LRR – leucine-rich repeat; Gly_zip – glycine zipper domain; SpoVK, EssC, EccE3-HerA – see text.

The first of the 3 variable regions flanked by the VWA domain-encoding genes includes an FtsK-like ATPase homologous to the Type VII secretion system factor EssC (Supplementary

Figure S15, Supplementary Table S1). EssC contains 3 tandem ATPase domains (D1, D2, and D3), with the Walker A/B motifs required for ATP hydrolysis present only in D1 and D2 [11, 12]. The CoCoNuT-associated homologs also possess 3 ATPase domains but differ in that only the central D2 domain homolog is predicted to be active. They are also distinguished from EssC by the presence of a coiled-coil that can exceed 200 residues in length and is fused at their N-terminus, whereas forkhead-associated domains and transmembrane helices are found in this position in EssC (Supplementary Table S1) [11, 12]. In addition, the region codes for two WXG100 proteins, one with the characteristic WxG motif and the other without (but with a similar predicted structure), as well as a coiled-coil fused to a restriction endonuclease-like domain with a D-ExK catalytic motif (Supplementary Figure S15, Supplementary Table S1) [13]. Finally, another protein similar to DNA mimics that bind the HU histone-like factor is encoded following the coiled-coil-nuclease fusion (Supplementary Figure S15, Supplementary Table S1) [14].

It appears likely that some or all of these factors are secreted, especially the WXG100 proteins, which are known to be secreted, with the prototypical example, ESAT-6, being a T-cell antigen diagnostic of *M. tuberculosis* infection [13]. These associated WXG100 proteins implicate the ATPases in defensive protein secretion, but as they lack transmembrane domains present in their EssC homologs, a different mechanism appears likely. The presence of coiled-coils at the N-termini of these proteins suggests that they might interact with the coiled-coils in the core CoCoNuT factors and/or with the associated coiled-coil-nuclease fusion. The FtsK superfamily ATPases form hexamers [95], so should such an interaction occur, they are likely compatible with the CnuB/McrB GTPase hexamer. A prior study that tangentially examined these operons in the context of the TerY-P triad pointed out that this WXG100/FtsK-like ATPase operon is likely a mobile element that can be found as a stand-alone secretion system in other genomes [15].

In the second of these Type III-A CoCoNuT extensions with two VWA domains, a SpoVK family of AAA+ ATPases homologous to p97/CDC48 is encoded adjacent to a protein containing a C-terminal bacteriocin-like glycine zipper motif (Supplementary Figure S15, Supplementary Table S1). CDC48 is involved in eukaryotic protein quality control, particularly the degradation of proteins synthesized from non-stop mRNA, where it is required to release nascent polypeptides from stalled ribosomes to enable proteolysis [16]. CDC48 contains two tandem ATPase domains, both of which are active; the CoCoNuT-associated homologs also contain tandem ATPases, but the Walker A/B motifs required to bind and hydrolyze ATP are conserved only in the C-terminal domain [17, 18]. As members of the AAA+ superfamily, these ATPases assemble into hexamers [18], similarly to the McrB GTPases. **A**t their C-termini, these ATPases are fused to domains of unknown function, namely, a leucine-rich repeat element and a beta-barrel domain structurally similar to biotin carrier proteins, suggesting that these proteins might be biotinylated (Supplementary Figure S15, Supplementary Table S1) [19]. The conserved association of the CDC48-like ATPases with these Type III-A CoCoNuTs, which encode the potentially tmRNA-binding SPB domain, seems to provide support for the scenario of tmRNA interaction. Structural analysis of the bacteriocin-like protein encoded in these loci indicates that it adopts an inactivated PD-(D/E)xK-type restriction endonuclease fold, potentially nucleic acid-binding (Supplementary Table S1). These proteins might be analogous to the restriction endonuclease-like factors in the FtsK/EssC homolog neighborhoods (Supplementary Figure S15).

The third variant of these extended Type III-A CoCoNuTs encodes a distinct member of the FtsK/HerA superfamily, which is also likely assembled into hexamers (Supplementary Figure S15, Supplementary Table S1) [20]. AF2 structural modeling suggests these enzymes are homologs of the ESX-3 Type VII secretion system factor EccE3 [21], albeit containing a unique beta-strand insertion of variable length. This EccE3-like domain is fused to an ATPase domain similar to the Type IV secretion system protein VirB4, which is involved in bacterial conjugation (Supplementary Figure S15, Supplementary Table S1) [22]. These systems also encode a small helical domain of unknown function in operonic association with the ATPase (Supplementary Figure S15). These genes might be involved in the mobilization of the locus via conjugation, or instead play a similar role in secretion as predicted for the EssC-like ATPases. However, WXG100 homologs, like those that strongly imply a secretion-related function for the EssC-like ATPases, are not encoded near these VirB4-like ATPases. Lastly, we observed that, unlike the EssC-like ATPases and the SpoVK-like proteins, these enzymes are not always encoded between VWA genes at the 5’ end of the operons, but in some cases, migrated to the 3’ end; in these cases, however, duplicated VWA domains are present at the 5’ end, a potential vestige of an extension that was recently lost or relocated.

Overall, these elaborations of Type III-A CoCoNuT systems resemble the TerY-P triads in that they could be stand-alone defensive cassettes that augment the effectiveness of the core CoCoNuT systems. It is unclear, however, why these types of factors are flanked by TerY-like VWA domains, as opposed to restriction systems such as Type I RM, GmrSD, and Druantia Type III, which are commonly associated with Type III-A CoCoNuTs as well, but are never so tightly integrated into the operon [23-25]. Type III-A systems embedded in these extended operons are annotated in the Supplementary Data.

While investigating this additional diversity of Type III-A CoCoNuTs, we observed that Type III-A CoCoNuTs in *Helicobacter* appeared to be translated using an alternate genetic code because gene predictions with the standard code divided the expected open reading frames into many small fragments. We were unable to identify a known alternative code that would yield the expected CoCoNuT gene products. Thus, a novel type of conditional or otherwise complex translation regulation likely occurs in these species, perhaps triggered by phage infection (for the accessions of identifiable Type III-A CoCoNuT factors in *Helicobacter*, see Supplementary Data).
